# Improved Glucose Homeostasis Following Vertical Sleeve Gastrectomy is Associated with Alternate Wiring of the Liver Molecular Clock in a Rat Model of Spontaneously-Occurring Type 2 Diabetes

**DOI:** 10.1101/2023.10.09.561476

**Authors:** Aurelie Le Lay, Francois Brial, Mark Lathrop, Christophe Magnan, Dominique Gauguier

## Abstract

Bariatric surgery is associated with remission of type 2 diabetes (T2D). We aimed to advance fundamental understanding of mechanisms involved in improved glucose homeostasis following vertical sleeve gastrectomy (VSG). We carried out a series of pathophysiological, behavioural and liver transcriptome analyses in lean rats of the Goto-Kakizaki (GK) model of polygenic T2D following VSG or sham operation. VSG and resulting sustained reduction in glucose intolerance were associated with significant changes in liver histology and lean mass, and nycthemeral feeding patterns and activity. Liver transcriptome profiling identified differentially regulated pathways between VSG and sham GK, including inflammatory and immune processes and fatty acid metabolism. Deeper analysis of the transcriptome dataset showed that expression of almost all main regulators of the molecular clock was significantly and co-ordinately affected by VSG. Comparisons with liver transcriptome data previously generated in GK and normoglycemic rats suggested that VSG results in a profound remodelling of the regulation of the molecular clock. Our findings shed light on relationships between the molecular clock and nycthemeral feeding and activity, which may contribute to long-term therapeutic consequences of VSG in the context of polygenic T2D in the absence of confounding effects of obesity.

## Introduction

Type 2 diabetes (T2D) and obesity are characterised by an etiology combining environmental influences and genetic risk factors (1). Bariatric surgery, which is routinely used in obese patients to induce sustained weight loss (2, 3), is also associated with improved glucose homeostasis, through mechanisms at least in part independent of weight loss and reduced caloric intake (4). Roux-en-Y gastric bypass (RYGB), vertical sleeve gastrectomy (VSG) and stomach sparing protocols (e.g. duodenal-jejunal and jejuno-ileal bypass, jejunectomy) have consistently demonstrated their beneficial role in T2D, but the biological processes involved remain poorly understood.

Preclinical models maintained in controlled conditions (diet, light/dark cycle) are powerful systems to investigate biological consequences of bariatric surgery (5). The inbred Goto-Kakizaki (GK) rat is a lean model of spontaneous T2D, obtained through genomic enrichment in naturally-occurring genetic polymorphisms over many generations of successive breeding of glucose intolerant outbred rats (6). Even though diabetes in the GK strain is caused by genes that have been localised in the rat genome by linkage analysis and functional genomics (Reviewed in (7))(8), bariatric surgery is able to improve glycemic control in this strain (9–11). We showed that reduction of hyperglycemia in gastrectomised GK rats is associated with alterations in gut microbiota architectural and bile acid metabolism (12).

To characterise the physiological and molecular mechanisms involved in diabetes remission following bariatric surgery, we carried out detailed analysis of nycthemeral feeding patterns and activity in gastrectomised GK rats and sham operated GK controls, followed by liver gene transcription profiling. Our findings suggest that mechanisms relevant to chronobiology are involved in VSG-promoted long term improvement in glucose homeostasis in this preclinical model of T2D in the absence of confounding effects of obesity.

## Material and Methods

### Animals

Inbred Goto-Kakizaki (GK/Ox) rats were bred in individually ventilated cages. Rats were maintained in a controlled environment of 12 h dark-light cycles, temperature (22–24°C) and relative humidity (50–60%). They had access to water and standard chow (SAFE, Augy, France) ad libitum. All procedures were performed in male rats under a licence (Ref. 5611 2016060311046952 v2) granted by the Charles Darwin Ethics Committee in Animal Experiment, Paris, France.

### Bariatric surgery in GK rats

Vertical sleeve gastrectomy (VSG) was performed in 14-week-old anesthetized male GK rats as previously described (12). Briefly, the lateral 80% of the stomach was excised to leave a tubular gastric remnant in continuity with the oesophagus, the pylorus and the duodenum. Control GK rats were sham operated through application of pressure with forceps along the oesophageal sphincter and the pylorus.

### Glucose tolerance tests

Intraperitoneal glucose tolerance tests (IPGTT) were performed in conscious rats one week before surgery and 4 weeks afterwards, when VSG-treated rats had recovered a body weight similar to that of sham-operated GK. Overnight fasted rats were injected intraperitoneally with a solution of glucose (1g/kg body weight). Blood samples were collected from the tail vein before glucose injection and 15, 30, 60, 120 and 180 minutes afterwards. Blood glucose was determined using an Accu-Check® Performa glucometer (Roche Diagnostics, Meylan, France). The increment of glucose values during the IPGTT was used to assess glucose tolerance.

### Nycthemeral feeding pattern recording and activity analysis

Three months after bariatric surgery, when body weight of VSG-treated GK rats returned to values of sham operated GK controls, spontaneous feeding and locomotor activity were measured using an automated system of feeding and activity recording (Labmaster, TSE Systems, Bad Homburg, Germany). Food consumption was recorded every 160 minutes over a period of three days. Diurnal and nocturnal activities were recorded using an infrared light beam-based locomotion monitoring system (beam breaks/h).

### Measurement of body mass composition

Fat mass and lean tissue mass were recorded before bariatric surgery and 26, 33, 52, 76 and 90 days after surgery using an Echo Medical Systems Echo MRI 100 (Whole Body Composition Analysers, EchoMRI, Houston, USA).

### Sample collection

At the end of the experiment, rats were fasted overnight and killed between 10am and 1pm by injection of sodium pentobarbital. Liver samples were harvested and either processed for histopathology analysis or snap-frozen in liquid nitrogen and stored at −80°C until mRNA preparation.

### Hepatic histopathology

Liver samples were snap frozen in OCT (VWR, Fontenay-sous-Bois, France) and 6-µm sections were prepared. Sections were rehydrated in PBS (Sigma-Aldrich, St Quentin, France) and stained with an Oil red O (ORO) solution (Sigma Aldrich, St Quentin, France). Slides were mounted with Vectashield (Eurobio Abcys, Les Ulis, France). For analysis of liver fibrosis, sections were stained with a rabbit anti-alpha smooth muscle actin (α-SMA) antibody (Abcam, ab124964, 1:500). A polymer-HRP detection system kit combined with DAB (Golden Bridge International, Blagnac, France) was used and slides were counter-stained with hematoxylin. Staining signals were quantified with and AxioScan microscope using the Visiopharm 2017.2 software.

### RNA preparation and RNA sequencing pipeline

Total RNA was extracted using the RNeasy RNA Mini Kit (Qiagen, Courtaboeuf, France). mRNA was fragmented and converted to cDNA, which was end-repaired, A-tailed and adapter-ligated before amplification and size selection. The prepared libraries were multiplexed and quality controlled before 51-nt paired end sequencing on an Illumina HiSeq2000 sequencer. RNA-Sequencing raw reads were processed using GenPipe (13). Briefly, reads were trimmed from the 3’ end to have a Phred score of at least 30 and filtered for a minimum length of 32 bp. Illumina sequencing adapters were clipped off from the reads using Trimmomatic v. 0.36 (14). Filtered reads were aligned to the rat genome reference (downloaded from UCSC) using STAR v. 2.5.3 with 2-passes mode (15). Read counts of Ensembl genes (version 84) were obtained using htseq-count v. 0.6.1 (16). Differential expression analyses at gene level were performed using DESeq2 (17).

### Biological pathway analysis

Gene Set Enrichment Analysis (GSEA) was performed for functional analysis of the liver transcriptomes (18). T-statistics of differential expression with default parameters and 1000 permutations were used. P-values were corrected using a significant level of 25% using the Benjamini-Hochberg (BH) method. The Kyoto Encyclopedia of Genes and Genomes (KEGG) (www.genome.jp/kegg), the Panther (www.pantherdb.org) and the Wikipathways (www.wikipathways.org) databases were used to detect enriched pathways in gastrectomised GK rats (19, 20). Over-representation analysis (ORA) was also performed. A significant level of 5% after BH correction for multiple testing of p-values was applied. Both GSEA and ORA analyses were performed using the R package WebGestaltR (version 3).

### Quantitative PCR

Quantitative RT-PCR analyses of candidate genes were performed using SYBR green assays (Life technologies, Saint Aubin, France). We used the housekeeping gene HPRT to normalize relative quantification of mRNA levels. Reactions were run on a qTower^3^ Real-Time PCR Thermal Cyclers (Analytik Jena). Oligonucleotide sequences are given in **Supplementary Table 1**.

## Results

### VSG improves glucose tolerance and liver histopathology in GK rats

We initially verified that VSG improves glucose homeostasis as previously reported in our GK colony (12). IPGTT were performed in the same animals before VSG or sham operation and 4 weeks afterwards. Sham operation did not affect the glycemic profile (**Figure 1a**) and the overall glucose response (**Figure 1b**) during the IPGTT. In contrast glucose tolerance was improved by VSG as reflected by the significant reduction of glycemia 120 and 180 minutes after glucose injection (**Figure 1c**), resulting in decreased cumulative glycemia (**Figure 1d**). Elevated hepatic triglycerides and anomalies in the regulation of liver lipid metabolism spontaneously occur in the GK rat (21). To assess the effect of VSG on liver lipid and fibrosis, we analysed liver sections of gastrectomised and sham operated GK. VSG induced a strongly significant reduction of liver fat content determined by Oil-Red-O staining of liver sections (**Figure 1e**). On the other hand, the hepatic level of α-SMA protein, a marker of fibrosis, was significantly more elevated in VSG rats than in sham operated controls, suggesting a residual inflammation following gastrectomy (**Figure 1f**).

**Figure 1.**
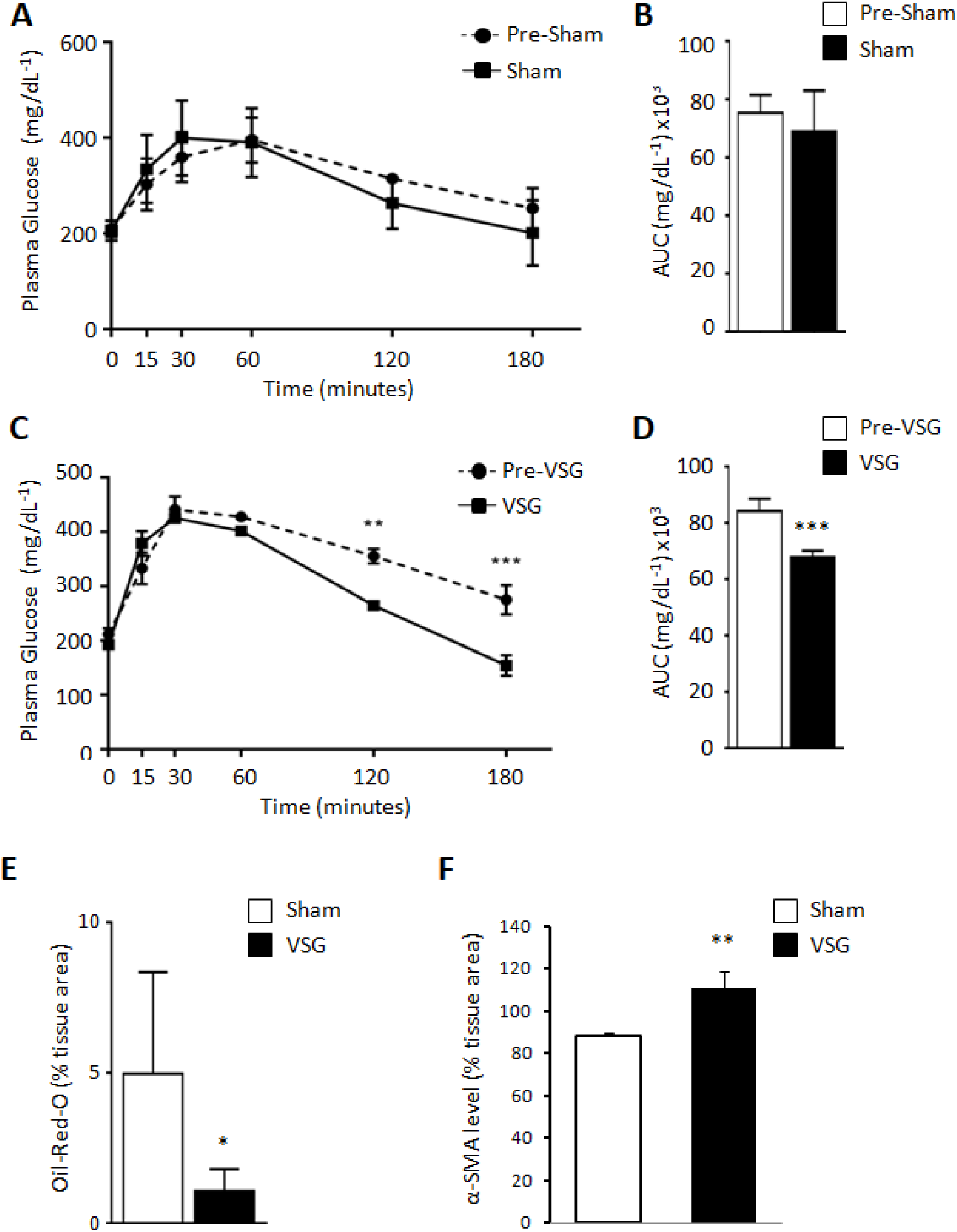
Effects of vertical sleeve gastrectomy (VSG) and sham operation on glucose tolerance and hepatic histopathology in GK rats. Intraperitoneal glucose tolerance tests (IPGTT) were carried out in the same animals before sham operation or VSG and 4 weeks afterwards (n=6 per group) (a-d). Liver sections prepared from sham operated and VSG GK rats 5 weeks after surgery were stained with Oil Red O (e) and anti-alpha smooth muscle actin (α-SMA) (f). Two sections of 4 rats per group were used for quantitative analysis. A Mann-Whitney test was used for statistical analysis. Results are means ± SEM. ***p<0.005, **p<0.01, *p<0.05 significantly different between VSG and sham operated animals.

### VSG results in changes in nycthemeral feeding patterns and activity in GK rats

Growing evidence supports a role of chrono-nutrition and temporal eating patterns in metabolic health (22). To test the hypothesis of an involvement of chronobiology in VSG-promoted improvement of glucose homeostasis, we performed nycthemeral recordings of food intake, which was repeated every 160 minutes over a period of three days, and activity in gastrectomised and sham operated GK rats. As expected sham-operated GK rats exhibited predominant nocturnal consumption of food (**Figure 2A**). In contrast, gastrectomised GK rats consumed significantly less food than sham operated controls during the dark phase, and showed an almost even consumption of food during the light and dark phases. These significant changes in nycthemeral feeding patterns in GK rats following VSG were associated with increased nocturnal activity (**Figure 2B**). The percentage of fat was similar in gastrectomised and sham-operated GK rats over the 90 days period following VSG (**Figure 2C**). In contrast, lean mass was significantly increased in VSG rats when compared to controls throughout the experiment (**Figure 2D**). These results support the involvement of mechanisms relevant to chronobiology in response to gastrectomy and improved glucose homeostasis.

**Figure 2.**
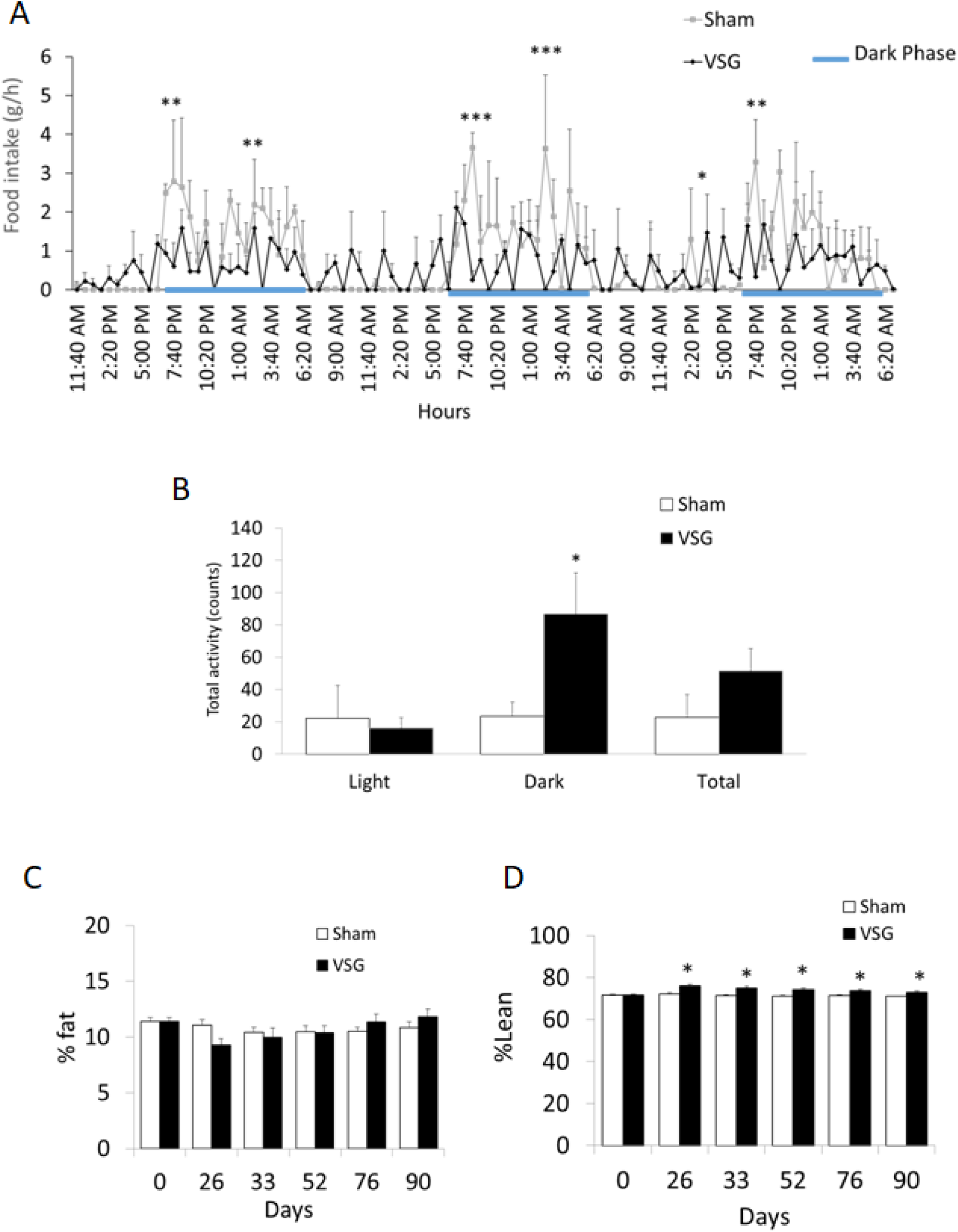
Nycthemeral feeding patterns and activity, and body composition following vertical sleeve gastrectomy (VSG) or sham operation in GK rats. Three months after bariatric surgery, spontaneous feeding was recorded every 160 minutes over a period of three days (A) and diurnal and nocturnal activities were determined (B). Fat mass (C) and lean tissue mass (D) were determined before surgery (Day 0) and 26, 33, 52, 76 and 90 days afterwards. Results are expressed as a percentage of body weight. Results are means ± SEM. ***p<0.005, **p<0.01, *p<0.05 significantly different between VSG and sham operated rats.

### Liver transcriptome is deeply altered following VSG and associated reduction of glucose intolerance in GK rats

To identify the molecular mechanisms underlying VSG-promoted improvement in glucose homeostasis, liver histopathology and nycthemeral behavior in the GK strain, we sequenced liver mRNA from gastrectomised and sham-operated GK rats. A total of 3102 genes were differentially expressed between the two rat groups, including 242 sequences corresponding to gene models, predicted genes and non coding RNA (**Supplementary Table 2**). There were equivalent numbers of down-regulated and up-regulated genes in response to VSG.

### Pathway analysis of liver transcriptome underlines the profound impact of VSG on multiple biological mechanisms

To identify biological functions affected by VSG and improved glucose homeostasis consecutive to VSG in the GK rat, we carried out gene set enrichment analysis (GSEA) of KEGG pathways. Among the 12,494 genes in our transcriptome dataset that were mapped to a unique EntrezGeneID, 5170 could be retrieved in KEGG and used for GSEA. A total of 33 biological pathways were significantly differentially enriched between VSG rats and sham operated controls (**Figure 3**). Normalised Enrichment Scores (NES) identified significantly up-regulated (n=19) or down-regulated (n=14) pathways (**Supplementary Table 3**). Details of genes contributing to the differential enrichment of these KEGG pathways in the rat models are given in **Supplementary Table 4.** The most significantly enriched pathways that differentiated VSG and sham operated GK rats were related to inflammatory and autoimmune processes, including “Type I diabetes mellitus” (NES=+1.851; P=0.004), “complement and coagulation cascades” (NES=+1.836; P=0.017), “primary immunodeficiency” (NES=+1.711; P=0.019), “natural killer cell mediated cytotoxicity” (NES=+1.708; P=0.019) and “chemokine signalling” (NES=+1.681; P=0.025) (**Figure 3, Supplementary Table 3**). In contrast, several metabolic pathways were significantly down-regulated in VSG GK rats, including “PPAR signalling” (NES=-2.074; P=0.004), “fatty acid metabolism” (NES=-1.818; P=0.016), “biosynthesis of unsaturated fatty acids” (NES=-1,799; P=0.013) and “peroxisome” (NES=-1.616; P=0.044).

**Figure 3.**
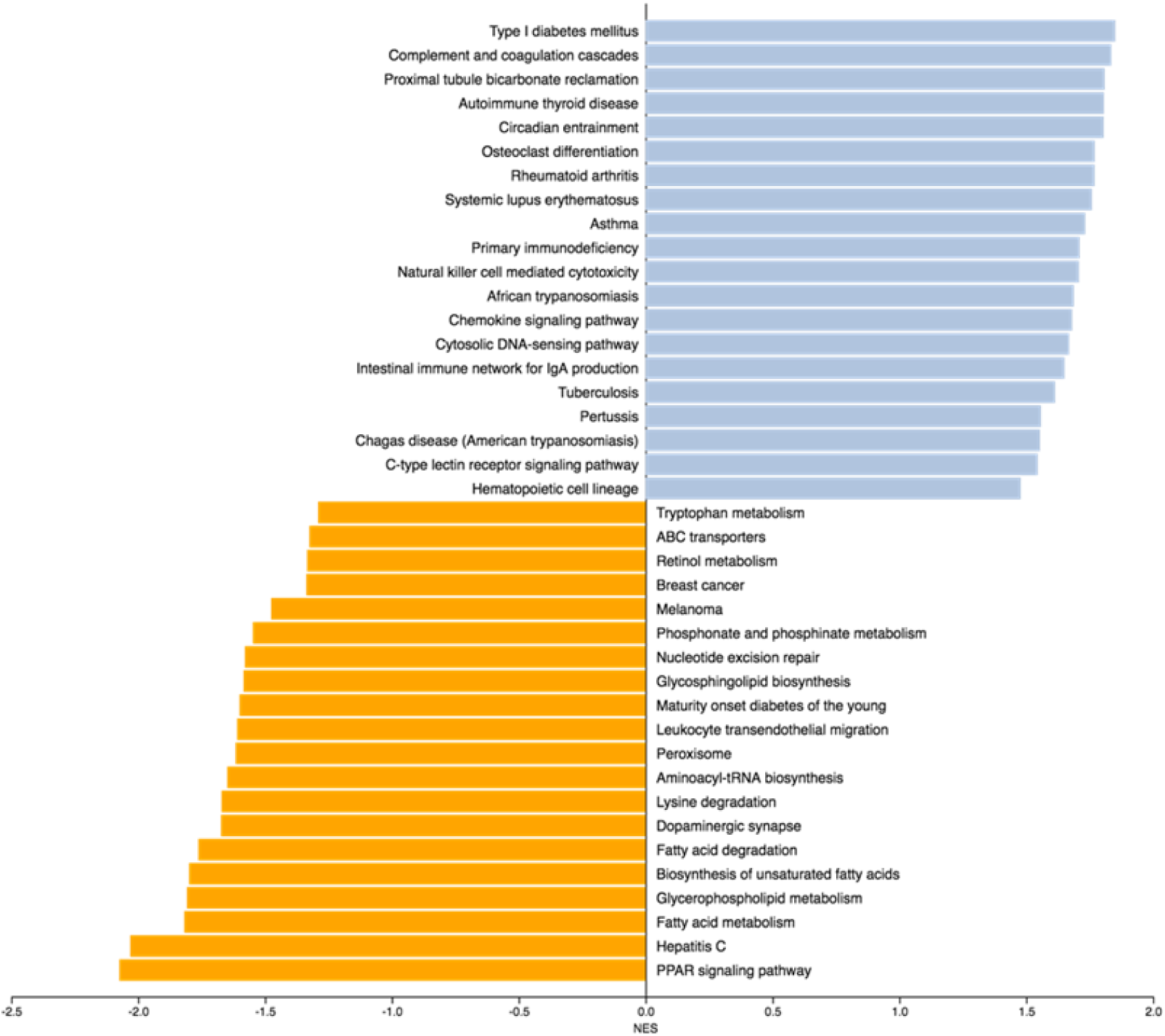
Pathways differentially enriched in GK rats following vertical sleeve gastrectomy (VSG) or sham operation. Gene set enrichment analysis (GSEA) of the liver transcriptomes of gastrectomised and sham operated GK rats was used to identify KEGG pathways up- or down-regulated, as illustrated by measures of a normalized enrichment score (NES). Blue bars indicate positive enrichment scores and orange bars indicate negative enrichment scores. RNA sequencing was performed in 4 rats per group. Details of statistics supporting pathway differential enrichment are given in **Supplementary Table 3**. Details of rno pathways can be found at www.genome.jp/kegg.

### Elements of the “PPAR signalling” pathway are down-regulated by VSG in the GK rat

The KEGG pathway PPAR signalling (rno03320) showed the strongest evidence of down-regulation in VSG GK rats, which may account for altered regulation of pathways related to fatty acid metabolism in this model (**Figure 2**). Expression of both core transcription factors in this pathway encoding the peroxisome proliferator activated receptor alpha (*Ppara*) and the retinoid X receptor gamma (*Rxrg*) was significantly decreased in VSG rats (Log2fold change (FC): −0.874; P=3.58×10^−5^ for *Ppara* and Log2FC: −0.505; P=0.003 for *Rxrg*) (**Table 1**). This is expected to have strong repercussions on the transcription of genes contributing to ketogenesis and fatty acid metabolism, which was a pathway (rno01212) significantly down-regulated in VSG rats. Significant alteration of the expression of the genes encoding 3-hydroxy-3-methylglutaryl-CoA synthase 2 (*Hmgcs2*) (Log2FC: −0.780; P=6.4×10^−4^), Acetyl-CoA C-acetyltransferase (*Acat2l1*) (Log2FC: +0.74, P=7.6×10^−4^), 3-oxoacid CoA-transferase (*Oxct1*) (Log2FC: +1.05, P=1.6×10^−3^), and both 3-hydroxybutyrate dehydrogenase 1 (*Bdh1*) (Log2FC: −0.87, P=7.5×10^−5^) and 2 (*Bdh2*) (Log2FC: +1.06, P=4.6×10^−4^) (**Supplementary Table 2**) suggests reduced ketogenesis in VSG rats. Down-regulated expression of genes involved in fatty acid oxidation (*Cpt1b, Cpt2*, *Acadl*, *Acadm*, *Acox1*), cholesterol metabolism (*Cyp8b1*), translocation of long-chain fatty acids across the plasma membrane (*Slc27a4*), fatty acid elongation (*Slc27a5*) and arachidonic acid metabolism (*Acsl4*, *Cyp4a1 Cyp4a2*) (**Table 1**) are also likely direct causes of *Ppara* and *Rxrg* decreased expression in VSG rats. Consistent with the reduction of liver fat in VSG rats, expression of three of the five perilipin genes was significant reduced (**Table 1**), including perilipin 2 (*Plin2*) which was massively down-regulated (Log2FC: −3.270; P=4.2×10^−194^).

**Table 1.**
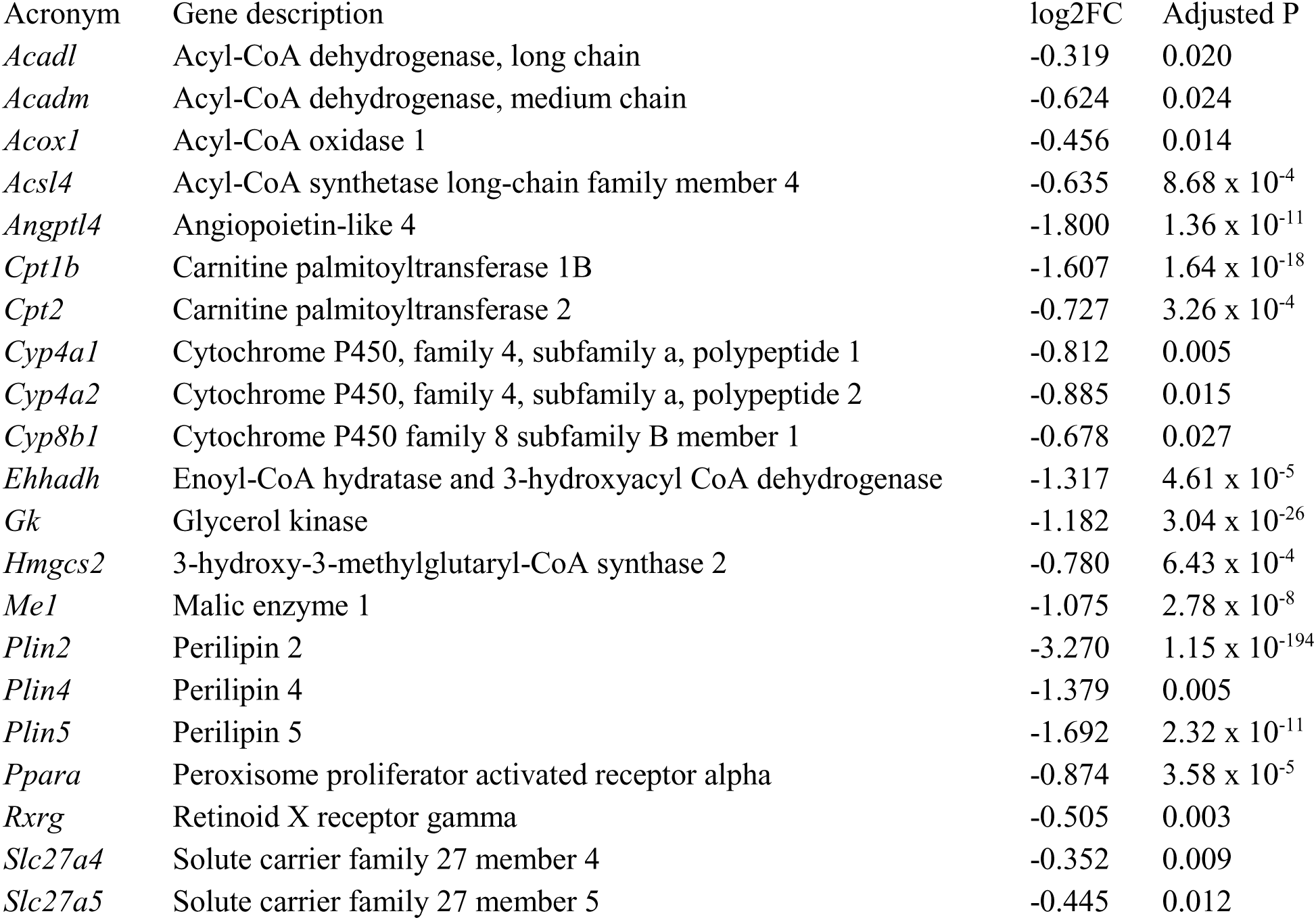
Genes contributing to down-regulated enrichment of the KEGG biological pathway “PPAR signaling” (rno03320) in gastrectomised GK rats. FC, fold change.

Altered regulation of inflammation and lipid metabolism were also identified when GSEA was carried out with Wikipathways (www.wikipathways.org) (**Supplementary Figure 3**) and when ORA was performed using the Wikipathways and the KEGG (**Supplementary Table 5**) and Reactome (**Supplementary Table 6**) databases. Interestingly, the latter pointed to overrepresentation of five pathways related to bile acid synthesis.

### Transcription regulation of key elements of the molecular clock is strongly altered by VSG

Analysis of transcriptome data through both GSEA and ORA of KEGG and Panther pathways consistently pointed to changes in the regulation of the molecular clock in VSG rats (**Supplementary Tables 3 and 5**). The KEGG pathway “circadian entrainment” (rno04713) was found significantly enriched through GSEA (NES=+1.805; P=0.021) (**Figure 3, Supplementary Table 3**). The pathway “circadian clock system” in the Panther database (P00015) was found significantly differentially regulated in the rat groups through both GSEA (NES=-0.9340, FDR=1.0) (**Supplementary Figure 1**) and ORA (P=0.0036; FDR=0.1404). ORA also indicated that the KEGG pathway “circadian rhythm” (rno04710) showed the strongest ratio of enrichment in the liver transcriptome of VSG rats (ratio: 3.02; p=0.0018) (**Supplementary Figure 2, Supplementary Table 5**).

The genes that contributed the most to enrichment of the KEGG pathway “circadian entrainment” were period circadian regulators *Per1* (Log2FC: +1.046; P=4.2×10^−8^), *Per2* (Log2FC: +2.000; P=1.9×10^−9^) and *Per3* (Log2FC: +3.793; P=3.0×10^−102^) (**Table 2**). The KEGG pathway “circadian rhythm” (rno04710), which was overrepresented in VSG rats, included these three genes and differentially expressed genes encoding the aryl hydrocarbon receptor nuclear translocator-like (*Arntl*, *Bmal1*) (Log2FC: −4.391; P=8.0×10^−105^), the neuronal PAS domain protein 2 (*Npas2*) (Log2FC: −3.647; P=4.7×10^−82^), the clock circadian regulator (*Clock*) (Log2FC: −1.298; P=3.3×10^−49^), the basic helix-loop-helix family, member e40 (*Dec1*, *Bhlhe40*) (Log2FC: +0.932; P=0.004) and member e41 (*Dec2*, *Bhlhe41*) (Log2FC: +2.839; P=2.4×10^−13^), the nuclear receptor subfamily 1, group D, member 1 (*Nr1d1, Rev-Erα*) (Log2FC: +1.352; P=3.3×10^−4^), cullin 1 (*Cul1*) (Log2FC: −0.223; P=0.021), the RAR related orphan receptor A (*Rora*) (Log2FC: −0.684; P=1.2×10^−3^), the S-phase kinase associated protein 1 (*Skp1*) (Log2FC: −0.222; P=0.041) and the cryptochrome circadian regulator 2 (*Cry2*) (Log2FC: +1.294; P=1.2×10^−7^) (**Supplementary Table 2**).

**Table 2.**
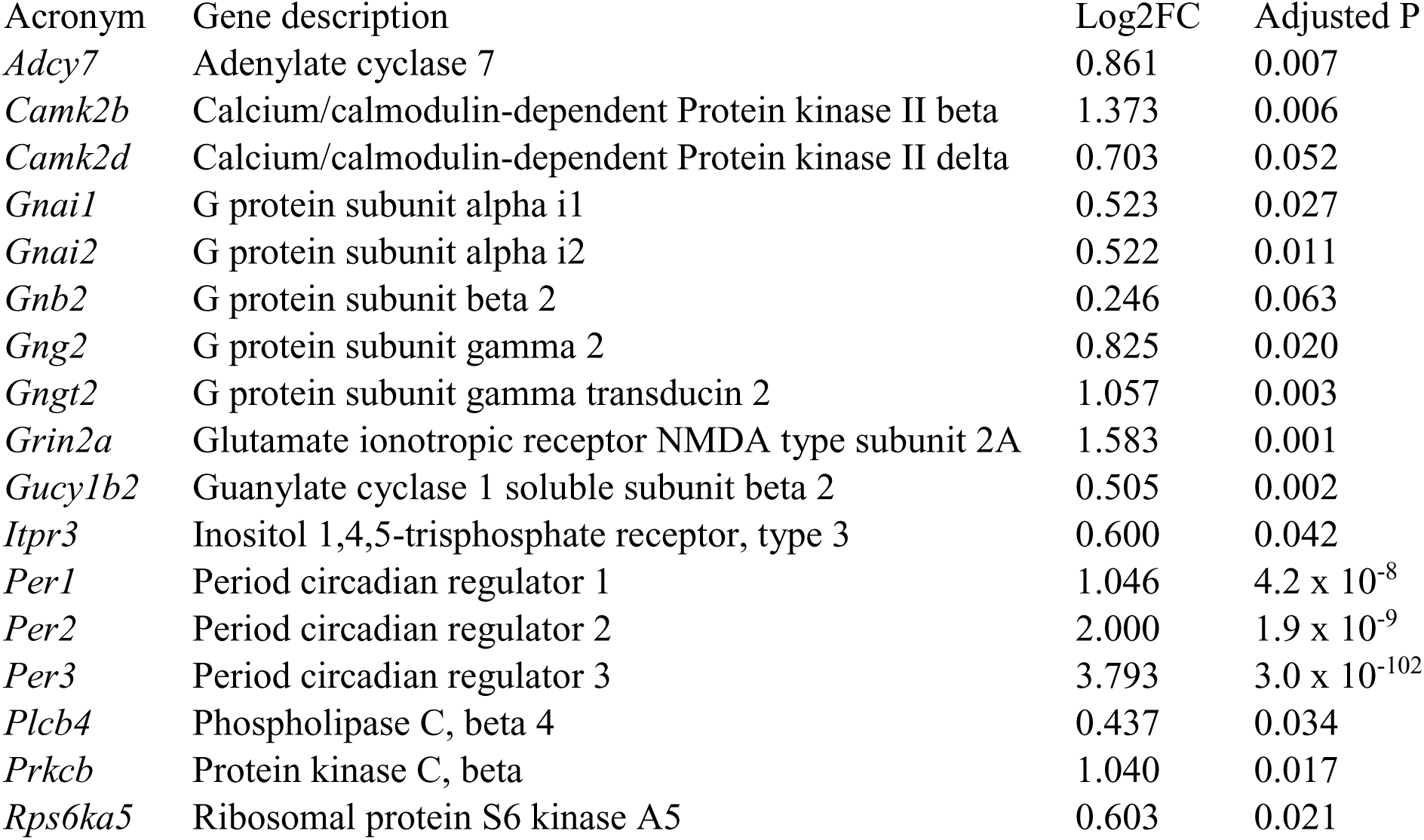
Genes contributing to the enrichment of the KEGG biological pathway “circadian entrainment” in gastrectomised GK rats. FC, fold change.

Deeper analyses identified altered expression of other elements of the molecular clock in VSG rats, including the D-box binding PAR bZIP transcription factor (*Dbp*), which was massively upregulated following VSG (Log2FC: +5.765; P=3.8×10^−216^), the circadian associated repressor of transcription (*Ciart*) (CHRONO) (Log2FC: +4.893; P=8.9 x 10^−46^), the nuclear receptor subfamily 1, group D, member 2 (*Nr1d2, Rev-Erβ*) (Log2FC: +1.291; P=8.0×10^−14^), the TEF transcription factor, PAR bZIP family member (*Tef*) (Log2FC: +1.382; P=5.7×10^−32^), timeless (*Tim*) (Log2FC: +0.644; P=0.004) and the ubiquitin specific peptidase 2 (*Usp2*) (Log2FC: +0.740; P=0.038) (**Supplementary Table 2**). Strikingly, 6 of the top 12 genes showing the strongest statistical evidence of differential expression in the transcriptome of VSG rats are central components of the molecular clock (*Dbp*, *Arntl*, *Per2*, *Npas2*, *Clock*, *Ciart*). Transcription of downstream targets of clock genes (eg. *Insig2*, *Elovl5*, *Acox1*, *Hadh*, *Hadha*, *Hadhb*, *Hnf4a*, *Hnf4g*, *Stat5a*, *Stat5b*) was down-regulated in VSG GK rats. These results indicate that coordinated and biologically coherent alteration in the expression of almost all known components of the hepatic molecular clock is a hallmark of the adaptation to VSG in a model of spontaneously-occurring T2D (**Figure 4**).

**Figure 4.**
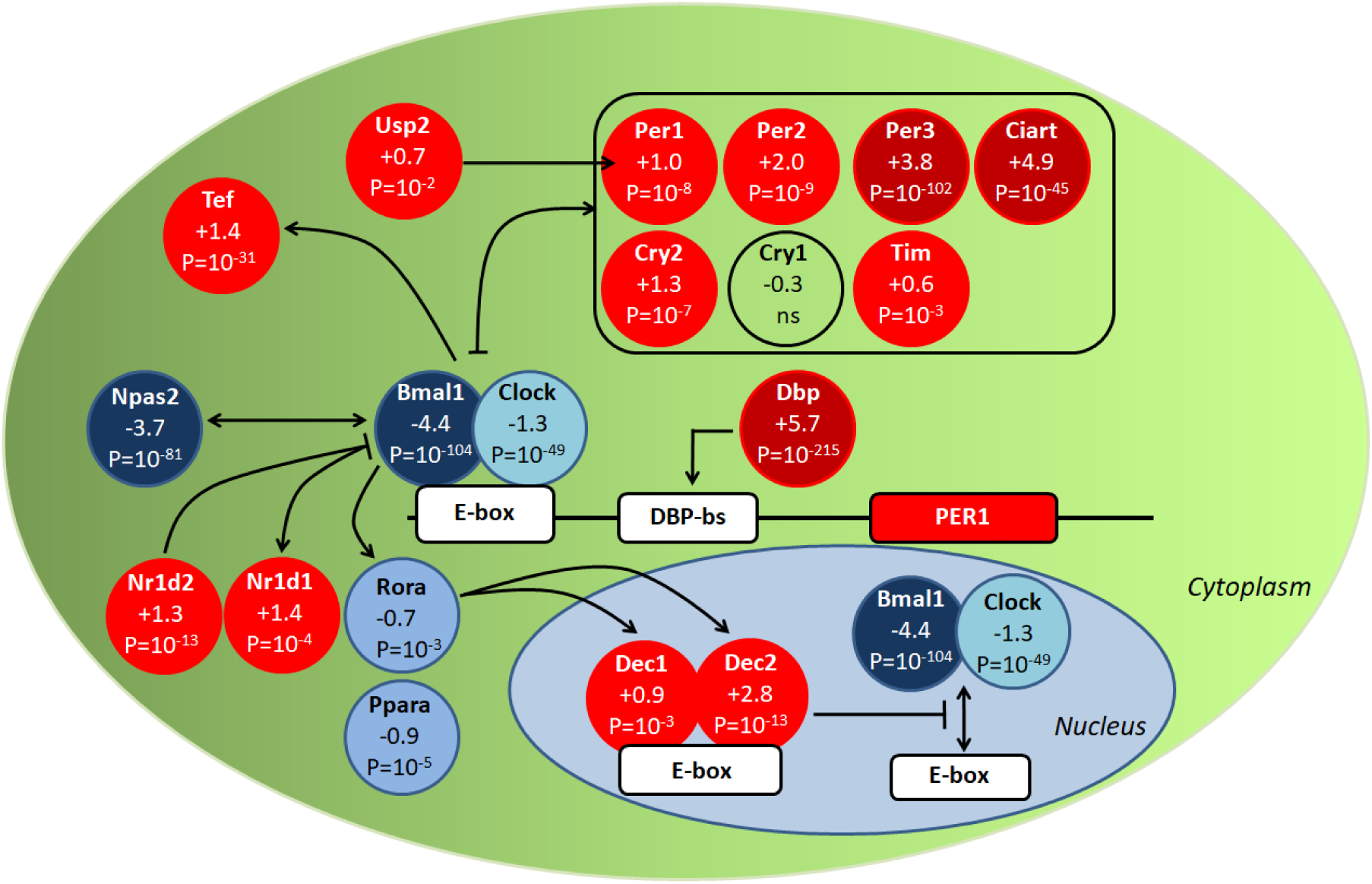
Schematic representation of liver transcription regulation of elements of the molecular clock in gastrectomised Goto-Kakizaki (GK) rats. Upregulated genes in gastrectomised rats are shown in red circles and down-regulated genes are shown in blue circles. Log2 Fold changes and corrected p-values are shown for each gene. Details of the genes are given in (**Supplementary Table 2**).

We confirmed by quantitative RT-PCR statistically significant differential expression of genes involved in inflammatory processes and the molecular clock (*Il1β*, *Il10*, *Ccl2*, *Arntl*, *Ciart*, *Cry2*, *Clock*, *Npas2*, *Per1*, *Per2*, *Per3* and *Rorα*) between VSG and sham operated GK rats (**Figure 5**).

**Figure 5.**
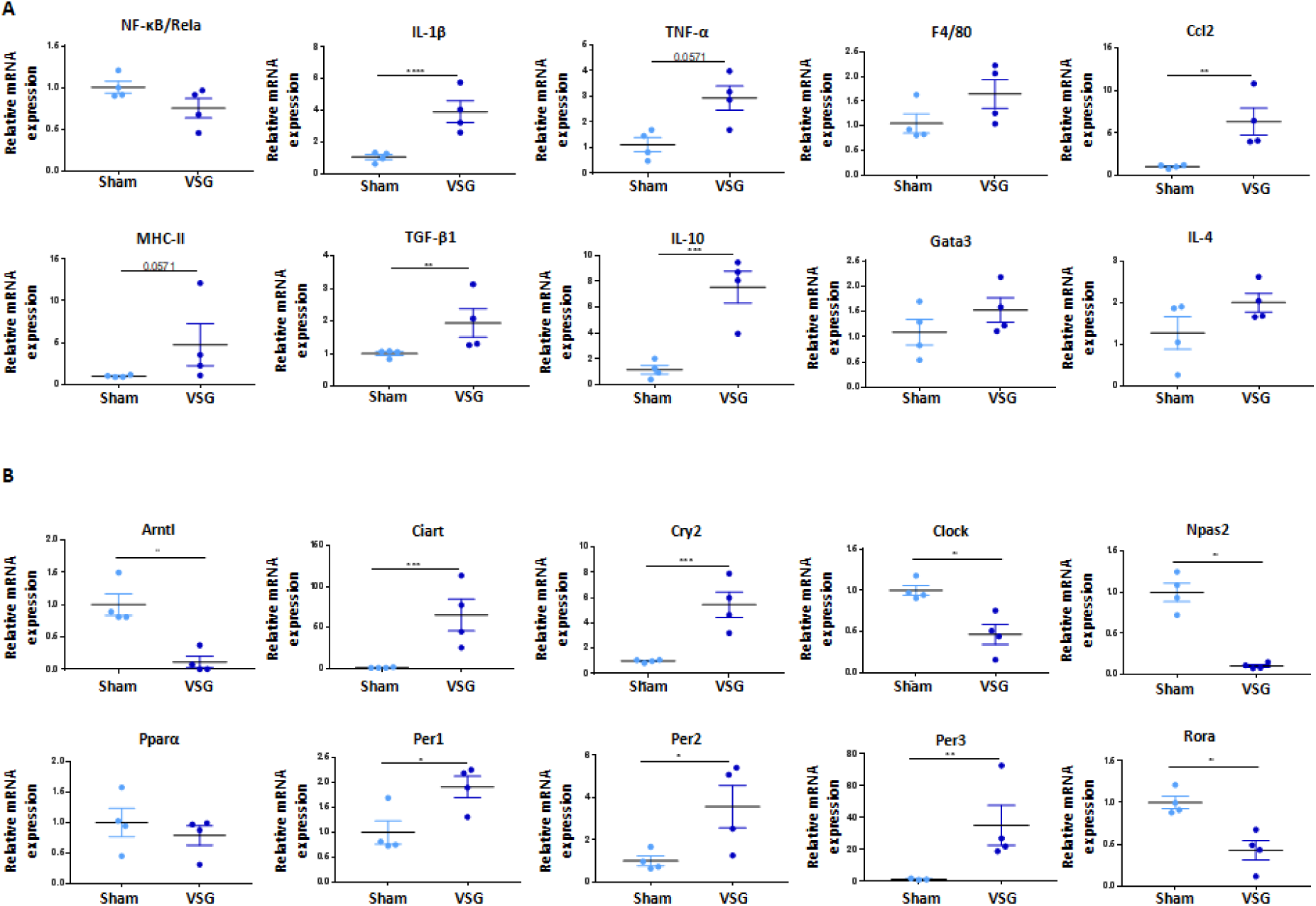
Expression of genes involved in inflammation (A) or circadian clock (B) in GK rats following vertical sleeve gastrectomy (VSG) or sham operation. Gene expression was determined analysed by quantitative PCR in liver of VSG and sham operated rats. Data are means ± SEM; n = 4 for each rat group; *p<0.05, **p<0.01, ***p<0.001, ****p<0.0001 significantly different between VSG and sham operated GK rats.

### Gene expression patterns suggest therapeutic consequences of VSG

To disentangle the possible pathophysiological relevance of VSG-promoted changes in liver gene expression and consecutive improvement in glucose homeostasis, we compared results from liver RNA sequencing in VSG and sham operated GK rats to those previously generated in GK/Ox and normoglycemic rats of the Brown Norway (BN) strain (23). Rats in the two studies were maintained in identical conditions and were killed in the same time window for organ collection. At the gene level, among the 3102 genes differentially expressed between gastrectomised GK and sham operated GK, only 567 (18%) were also differentially expressed between GK and BN (**Supplementary Table 7**). Of note, as many as 437 of these genes (77%) showed opposite expression patterns (**Figure 6**).

**Figure 6.**
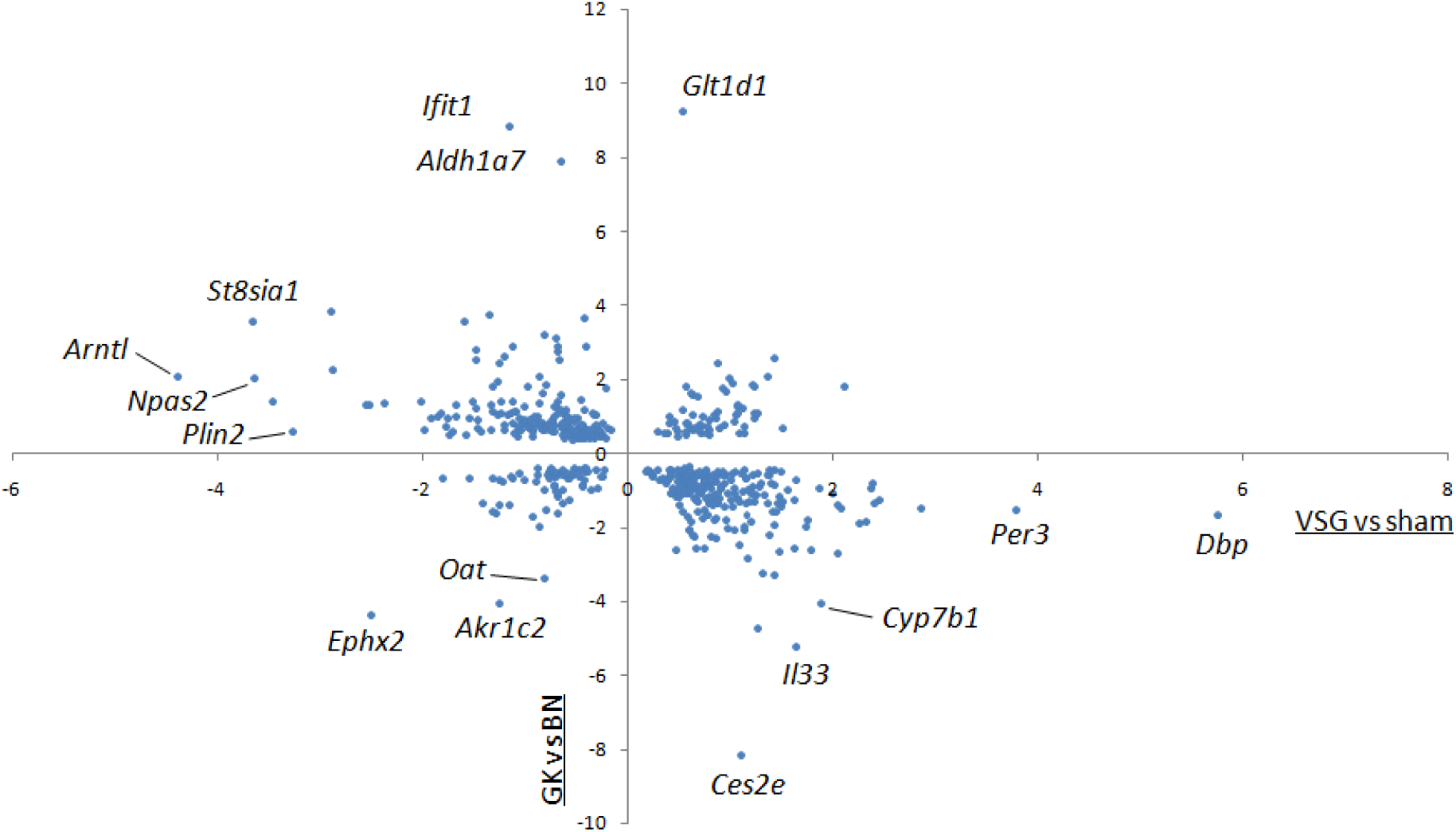
Comparative analysis of differentially expressed genes between gastrectomised Goto-Kakizaki (GK) rats and sham-operated GK rats and between GK and normoglycemic rats of the Brown Norway strain. Normalised enrichment scores (NES) of the 567 genes significantly differentially expressed in liver in the two comparisons are plotted to illustrate divergent and conserved liver gene expression patterns. Transcriptome (RNA sequencing) data in GK and Brown Norway rats are from Kaisaki et al (23). Details of the statistics and gene description are given in **Supplementary Table 7**.

The profound remodeling of gene transcription regulation following VSG was also suggested by results from biological pathway analysis. GSEA carried out in the two datasets identified only seven overlapping KEGG pathways in the two comparisons (“asthma”, “autoimmune thyroid disease”, “biosynthesis of unsaturated fatty acids”, “fatty acid degradation”, “peroxisome”, “PPAR signaling” and “type I diabetes mellitus”). However, the genes contributing to enrichment of these pathways were largely different in the two datasets. For example, only six genes of the fatty acid degradation pathway (*Acadm*, *Cyp4a1*, *Cyp4a2*, *Eci1*, *Ehhadh*, *Hadha*) and the PPAR signalling pathway (*Acadm*, *Angptl4*, *Cyp4a1*, *Cyp4a2*, *Cyp8b1*, *Ehhadh*) showed concordant differential expression and direction of expression change (**Supplementary Table 7**). On the other hand, expression of *Plin2* in the latter pathway showed contrasting direction of changes in the two datasets, suggesting that VSG contributes to reduce liver fat content in the GK. Similarly, the six genes in the “type I diabetes mellitus” pathway that were consistently differentially expressed in the two comparisons (*Il1a*, *Ptprn*, *Rt1Ba*, *Rt1Bb*, *Rt1Da*, *Rt1Db1*) showed opposite direction of transcription change (**Supplementary Table 7**).

At the gene level, VSG-promoted alternative regulation in gene expression patterns following VSG in GK rats was particularly significant for several genes involved in the molecular clock (*Arntl*, *Dbp*, *Npas2*, *Per1*, *Per2*, *Per3*, *Tef*, *Tim* and *Usp2*) (**Figure 5**, **Supplementary Table 7)**. The remaining genes of the molecular clock differentially expressed between gastrectomised GK and sham operated GK did not show evidence of significant differential expression between GK and BN rats.

Collectively, the profound remodeling of hepatic gene expression and evidence of alternative wiring of the liver molecular clock in response to VSG may contribute to the therapeutic consequences of bariatric surgery in the GK strain through nycthemeral alterations in feeding patterns and activity, and underline the impact of chronobiology in T2D etiopathogenesis.

## Discussion

We report evidence of alterations in chronobiology and hepatic gene expression occurring three months after VSG in the GK rat. VSG-promoted changes in nycthemeral feeding patterns and activity and in molecular pathways linked to the circadian clock, inflammation and lipid metabolism underlie responses to improved glucose homeostasis and reduced liver fat following gastrectomy. These results shed light on mechanisms associated with surgically-induced improvements in glucose homeostasis in a model characterized by spontaneously-occurring glucose intolerance in the absence of confounding effects of obesity.

We provide confirmatory evidence of the beneficial effects of bariatric surgery on diabetes, which was previously demonstrated in the GK strain (10–12, 24–27), in Otsuka Long-Evans Tokushima Fatty (OLETF) rats (28) and in streptozotocin-treated rats (10). Reduction in liver fat, inhibition of hepatic expression of *Plin2* and changes in bile acid synthesis in gastrectomised GK rats in our study support reports of improved fatty liver disease (29) and increased bile acid metabolism (30) following bariatric surgery. Activation of the farnesoid X receptor (FXR, NR1H4) and down-regulated expression of cholesterol 7α-hydroxylase (CYP7A1) account for the beneficial effects of bariatric surgery on glucose homeostasis (31, 32). Counterintuitive down-regulated expression of *Fxr* (Log2FC: −0.378; P=0.016) and upregulated expression of *Cyp7a1* (Log2FC: +1.440; P=1.2×10^−5^) in liver of gastrectomised GK rats in our study agree with improved hepatic steatosis and insulin sensitivity and increased bile acid metabolism in mice overexpressing *Cyp7a1* (33). These mice also exhibit decreased expression of *Cyp8b1*, which we also detected in VSG GK rats (Log2FC: −0.678; P=0.027), and contribute to increase the pool of bile acids as demonstrated in *Cyp8b1* knockout mice (34). These data suggest that stimulation of bile acid synthesis may occur in VSG GK rats through coordinated changes in the expression of *Cyp7a1* and *Cyp8b1*.

Transcriptome data in VSG GK rats suggest decrease in lipid-related pathways and reduction of fatty liver through down-regulated expression of both *Plin2* and *Plin5*. This is supported by reports of protection against fatty liver disease following inactivation of *Plin2* (35) and *Plin5* (36). Comparative analysis of liver transcriptome data in VSG and sham-operated GK rats and in GK and rats of the normoglycemic Wistar (21) and BN (23) strains was used to reveal transcriptional changes accounting for the therapeutic impact of gastrectomy in the GK. We noted discrepant directions of gene expression changes in the two comparisons for many genes encoding proteins involved in lipogenesis (*Acly*, *Fas*, *G6pd*, *Me1*) and fatty acid modification (*Elovl5*, *Scd1*), trafficking (*Fatp4*) and degradation (*Acox1*, *Acacb*), which indicate extensive changes in lipid metabolism following VSG. Consistent elevation of *Pparα* expression in GK rats when compared to Wistar (21) and BN (23) rats and contrasting down-regulation of its expression in VSG GK rats is one of the most biologically significant transcriptional adaptation to VSG accounting for altered regulation of lipid metabolism. Such strong divergences in hepatic transcription regulations in these model comparisons indicate an extensive remodelling of gene expression in response to VSG, which may result in beneficial physiological effects. Data from feeding and activity recordings in gastrectomised and sham operated GK rats suggest that VSG in diabetic individuals results in modulating chronobiology and chrono-nutrition characterised by adjustment of nycthemeral feeding pattern and increased nocturnal activity. There is increasing interest in the contribution of modified circadian rhythms in the etiopathogenesis of T2D and obesity (37). Modern-day lifestyle has altered circadian biology and dietary patterns, resulting in disruption of feeding–fasting cycle and metabolic processes (38, 39). Relationships between VSG, meal frequency and the circadian clock were recently addressed in mice (40). Time-restricted feeding has beneficial effects on metabolic health and glycemic control (41, 42), but the direct involvement of the circadian clock in rhythmic food intake through expression of core clock genes remains debated (43, 44).

Coordinated changes in the expression of the vast majority of genes involved in the hepatic molecular clock was one of the most remarkable transcriptional consequences of VSG in GK rats, which concurs with altered feeding patterns and activity in these rats. Core clock genes synchronise transcriptional and translational feedback loops and regulate metabolism in health and disease (45, 46). We observed increased expression of *Dbp*, *Per1*, *Per2* and *Per3* and decreased expression of *Naps2*, *Bmal1* and *Clock*, which fit the known patterns of feedback loop of transcription regulation of the pathway. The strongest and most significant upregulated gene in response to VSG in the GK rat was *Dbp*, which was proposed as an upstream regulator of the clock system (47). Of note, expression of genes regulating nicotinamide metabolism (*Nampt* and *Nmrk1*), which are rhythmically expressed in liver and involved in the autonomous hepatic clock (48), was upregulated in VSG rats. Strongly discordant expression patterns of central elements of the molecular clock (*Arntl*, *Dbp*, *Npas1*, *Per1*, *Per2*, *Per3*, *Tef*, *Usp2*) between VSG GK rats and sham-operated GK controls, and between GK and normoglycemic BN rats (23), (**supplementary Figure 4**) suggests that the therapeutic contribution of VSG in GK rats involves alternative wiring of the liver clock transcriptome.

## Conclusions

Our findings underline the breadth of behavioural and molecular mechanisms that collectively contribute to improved glycemic control consecutive to bariatric surgery in a model of T2D devoid of obesity. Causal relationships between physiological improvements in response to bariatric surgery, altered transcription regulation of the molecular clock, nycthemeral feeding patterns and activity, as well as changes in gut microbiota architecture and bile acid metabolism that we previously identified in gastrectomised GK rats (12), remain to be disentangled. Evidence of regulation of *Cyp7a1* and *Insig2* by *Nr1d1* (49) and binding of DBP to the promoters of *Cyp7a1* (50) and *Per2* (51), suggest connections between bile acid metabolism and the circadian clock in VSG GK rats. Our data contribute to enhanced knowledge in fundamental mechanisms contributing to the therapeutic outcomes of bariatric surgery in T2D and underline the importance of chronobiology in the regulation of glucose homeostasis.

## Authors contribution

M.L., C.M., E.G. and D.G. conceived the project. A.L.L, M.V. and C.R. performed sleeve surgery in GK rats. A.L.L. and F.B. performed animal procedures, RNA sequencing and qPCR analysis. X.S. analysed RNA sequencing data. M.R and K.A. carried out biological pathway analyses. D.G. wrote the manuscript.

## Abbreviations

BN: Brown Norway
GK: Goto-Kakizaki
GSEA: Gene Set Enrichment Analysis
IPGTT: Intraperitoneal Glucose Tolerance Test
KEGG: Kyoto Encyclopedia of Genes and Genomes
NES: Normalised Enrichment Scores
OLETF: Otsuka Long-Evans Tokushima Fatty
ORA: Over-Representation Analysis
P.copri: Prevotella copri
RYGB: Roux-en-Y gastric bypass
VSG: Vertical Sleeve Gastrectomy

