## Supplementary Table 4 for "Improved Glucose Homeostasis Following Vertical Sleeve Gastrectomy is Associated with Alternate Wiring of the Liver Molecular Clock in a Rat Model of Spontaneously-Occurring Type 2 Diabetes"

**Supplementary Table 4. Genes contributing to the differential enrichment of KEGG biological pathways in gastrectomised and sham operated Goto-Kakizaki (GK) rats.** Gene set enrichment analysis (GSEA) of the liver transcriptomes of gastrectomised and sham operated GK rats was used to identify up- or down-regulated KEGG pathways. FC, fold change.

| **Type I diabetes mellitus** | | | |  |  |  |
| --- | --- | --- | --- | --- | --- | --- |
| **Acronym** | | **Gene description** | | **ENSEMBL Reference** | **log2FC** | **Adjusted P** |
| Cd28 | | CD28 molecule | | ENSRNOG00000010283 | 0.787 | 0.15200581 |
| Cd80 | | CD80 molecule | | ENSRNOG00000001527 | 1.409 | 0.00152716 |
| Cd86 | | CD86 molecule | | ENSRNOG00000038835 | 0.950 | 0.00354688 |
| Fas | | Fas cell surface death receptor | | ENSRNOG00000019142 | 1.155 | 0.00142758 |
| Gzmb | | granzyme B | | ENSRNOG00000049976 | 1.016 | 0.02904778 |
| Il1a | | interleukin 1 alpha | | ENSRNOG00000004575 | 1.152 | 0.0002448 |
| Il1b | | interleukin 1 beta | | ENSRNOG00000004649 | 1.714 | 2.58E-05 |
| Prf1 | | Perforin 1 | | ENSRNOG00000000562 | 0.571 | 0.09305383 |
| Ptprn | | protein tyrosine phosphatase, receptor type, N | | ENSRNOG00000019587 | 1.324 | 0.00102865 |
| RT1-A1 | | RT1 class Ia, locus A1 | | ENSRNOG00000038999 | 0.230 | 0.08979376 |
| RT1-Ba | | RT1 class II, locus Ba | | ENSRNOG00000000451 | 1.408 | 0.00402879 |
| RT1-Bb | | RT1 class II, locus Bb | | ENSRNOG00000032708 | 1.374 | 0.00139534 |
| RT1-CE4 | | RT1 class I, locus CE4 | | ENSRNOG00000039744 | 0.236 | 0.08789066 |
| RT1-CE5 | | RT1 class I, locus CE5 | | ENSRNOG00000000723 | 0.659 | 0.07770075 |
| RT1-Da | | RT1 class II, locus Da | | ENSRNOG00000032844 | 1.307 | 0.00820569 |
| RT1-Db1 | | RT1 class II, locus Db1 | | ENSRNOG00000033215 | 1.395 | 0.00422669 |
| RT1-N3 | | RT1 class Ib, locus N3 | | ENSRNOG00000000795 | 0.464 | 0.15287154 |
| **Complement and coagulation cascades** | | | | | |  |
| **Acronym** | **Gene description** | | | **ENSEMBL Reference** | **log2FC** | **Adjusted P** |
| Bdkrb2 | bradykinin receptor B2 | | | ENSRNOG00000047300 | 0.846 | 0.12133858 |
| C1qa | complement C1q A chain | | | ENSRNOG00000012807 | 0.642 | 0.06250174 |
| C1qb | complement C1q B chain | | | ENSRNOG00000012749 | 0.770 | 0.02624876 |
| C1qc | complement C1q C chain | | | ENSRNOG00000012804 | 0.725 | 0.04279283 |
| C1r | complement C1r | | | ENSRNOG00000011796 | 0.516 | 0.11635859 |
| C1s | complement C1s | | | ENSRNOG00000011971 | 0.691 | 0.0256251 |
| C3 | complement C3 | | | ENSRNOG00000046834 | 0.777 | 0.01474009 |
| C3ar1 | complement C3a receptor 1 | | | ENSRNOG00000009211 | 0.920 | 0.03341037 |
| C4a | complement C4A | | | ENSRNOG00000047657 | 0.121 | 0.83302637 |
| C4bpa | complement component 4 binding protein, alpha | | | ENSRNOG00000004062 | 0.821 | 0.00996253 |
| C4bpb | complement component 4 binding protein, beta | | | ENSRNOG00000004125 | 0.934 | 4.78E-05 |
| C5ar1 | complement C5a receptor 1 | | | ENSRNOG00000047800 | 0.940 | 0.02306636 |
| C6 | complement C6 | | | ENSRNOG00000024115 | 0.942 | 0.02349783 |
| C9 | complement C9 | | | ENSRNOG00000013736 | 1.047 | 0.01328089 |
| Cd55 | CD55 molecule (Cromer blood group) | | | ENSRNOG00000003927 | 0.802 | 0.11374796 |
| Cfd | complement factor D | | | ENSRNOG00000033564 | 0.682 | 0.05164728 |
| Cfh | complement factor H | | | ENSRNOG00000030715 | 0.685 | 0.04170008 |
| Cfi | complement factor I | | | ENSRNOG00000053400 | 0.217 | 0.06455537 |
| Cpb2 | carboxypeptidase B2 | | | ENSRNOG00000010935 | 0.420 | 0.03100581 |
| F2r | Coagulation factor II (thrombin) receptor | | | ENSRNOG00000048043 | 0.747 | 6.77E-05 |
| F2rl2 | coagulation factor II (thrombin) receptor-like 2 | | | ENSRNOG00000018054 | 0.934 | 0.07375596 |
| F7 | coagulation factor VII | | | ENSRNOG00000032737 | 0.477 | 0.03199765 |
| Itgax | integrin subunit alpha X | | | ENSRNOG00000036703 | 1.078 | 1.69E-05 |
| Itgb2 | integrin subunit beta 2 | | | ENSRNOG00000001224 | 0.903 | 0.01454084 |
| Klkb1 | kallikrein B1 | | | ENSRNOG00000014118 | 0.515 | 0.00211067 |
| Mbl1 | mannose-binding lectin (protein A) 1 | | | ENSRNOG00000011706 | 0.336 | 0.02717533 |
| Plau | plasminogen activator, urokinase | | | ENSRNOG00000010516 | 0.839 | 0.02738359 |
| Plaur | plasminogen activator, urokinase receptor | | | ENSRNOG00000037931 | 1.052 | 0.00359474 |
| Procr | protein C receptor | | | ENSRNOG00000019330 | 0.612 | 0.06087049 |
| Serpine1 | serpin family E member 1 | | | ENSRNOG00000001414 | 1.898 | 1.20E-18 |
| Serping1 | serpin family G member 1 | | | ENSRNOG00000007457 | 0.601 | 0.06757624 |
| Tfpi | tissue factor pathway inhibitor | | | ENSRNOG00000005039 | 0.421 | 0.00257171 |
| Thbd | thrombomodulin | | | ENSRNOG00000004687 | 0.443 | 0.09110132 |
| Vsig4 | V-set and immunoglobulin domain containing 4 | | | ENSRNOG00000038132 | 0.633 | 0.06625092 |
| Vwf | von Willebrand factor | | | ENSRNOG00000019689 | 0.870 | 0.00049617 |
| **Autoimmune thyroid disease** | | | | |  |  |
| **Acronym** | **Gene description** | | | **ENSEMBL Reference** | **log2FC** | **Adjusted P** |
| Cd28 | CD28 molecule | | | ENSRNOG00000010283 | 0.787 | 0.15200581 |
| Cd40 | CD40 molecule | | | ENSRNOG00000018488 | 0.916 | 0.0160465 |
| Cd80 | CD80 molecule | | | ENSRNOG00000001527 | 1.409 | 0.00152716 |
| Cd86 | CD86 molecule | | | ENSRNOG00000038835 | 0.950 | 0.00354688 |
| Fas | Fas cell surface death receptor | | | ENSRNOG00000019142 | 1.155 | 0.00142758 |
| Gzmb | granzyme B | | | ENSRNOG00000049976 | 1.016 | 0.02904778 |
| Il10 | interleukin 10 | | | ENSRNOG00000004647 | 1.552 | 6.23E-05 |
| Prf1 | Perforin 1 | | | ENSRNOG00000000562 | 0.571 | 0.09305383 |
| RT1-A1 | RT1 class Ia, locus A1 | | | ENSRNOG00000038999 | 0.230 | 0.08979376 |
| RT1-Ba | RT1 class II, locus Ba | | | ENSRNOG00000000451 | 1.408 | 0.00402879 |
| RT1-Bb | RT1 class II, locus Bb | | | ENSRNOG00000032708 | 1.374 | 0.00139534 |
| RT1-CE4 | RT1 class I, locus CE4 | | | ENSRNOG00000039744 | 0.236 | 0.08789066 |
| RT1-CE5 | RT1 class I, locus CE5 | | | ENSRNOG00000000723 | 0.659 | 0.07770075 |
| RT1-Da | RT1 class II, locus Da | | | ENSRNOG00000032844 | 1.307 | 0.00820569 |
| RT1-Db1 | RT1 class II, locus Db1 | | | ENSRNOG00000033215 | 1.395 | 0.00422669 |
| RT1-N3 | RT1 class Ib, locus N3 | | | ENSRNOG00000000795 | 0.464 | 0.15287154 |
| **Osteoclast differentiation** | | | | |  |  |
| **Acronym** | **Gene description** | | | **ENSEMBL Reference** | **log2FC** | **Adjusted P** |
| Acp5 | acid phosphatase 5, tartrate resistant | | | ENSRNOG00000046261 | 0.476 | 0.02608932 |
| Blnk | B-cell linker | | | ENSRNOG00000013967 | 0.785 | 0.04865425 |
| Btk | Bruton tyrosine kinase | | | ENSRNOG00000052407 | 0.890 | 0.03487453 |
| Csf1 | colony stimulating factor 1 | | | ENSRNOG00000018659 | 0.810 | 0.02787173 |
| Csf1r | colony stimulating factor 1 receptor | | | ENSRNOG00000018414 | 0.930 | 0.03072439 |
| Ctsk | cathepsin K | | | ENSRNOG00000021155 | 0.490 | 0.01265361 |
| Cyba | cytochrome b-245 alpha chain | | | ENSRNOG00000013014 | 1.114 | 0.00195734 |
| Fcgr1a | Fc fragment of IgG receptor Ia | | | ENSRNOG00000021199 | 1.110 | 0.00046746 |
| Fcgr3a | Fc fragment of IgG receptor IIIa | | | ENSRNOG00000024382 | 1.197 | 0.00522427 |
| Fosl2 | FOS like 2, AP-1 transcription factor subunit | | | ENSRNOG00000052357 | 0.577 | 0.09305383 |
| Fyn | FYN proto-oncogene, Src family tyrosine kinase | | | ENSRNOG00000000596 | 0.849 | 0.00048507 |
| Gab2 | GRB2-associated binding protein 2 | | | ENSRNOG00000011882 | 0.613 | 0.0808639 |
| Ifngr1 | interferon gamma receptor 1 | | | ENSRNOG00000012074 | 0.658 | 0.00243399 |
| Ikbkb | inhibitor of nuclear factor kappa B kinase subunit beta | | | ENSRNOG00000019073 | 0.374 | 0.08821073 |
| Il1a | interleukin 1 alpha | | | ENSRNOG00000004575 | 1.152 | 0.0002448 |
| Il1b | interleukin 1 beta | | | ENSRNOG00000004649 | 1.714 | 2.58E-05 |
| Il1r1 | interleukin 1 receptor type 1 | | | ENSRNOG00000014504 | 1.170 | 0.0023331 |
| Lck | LCK proto-oncogene, Src family tyrosine kina | | | ENSRNOG00000009705 | 0.807 | 0.00349529 |
| Lcp2 | lymphocyte cytosolic protein 2 | | | ENSRNOG00000005620 | 0.798 | 0.02527559 |
| Lilrb3 | leukocyte immunoglobulin like receptor B3 | | | ENSRNOG00000046683 | 0.953 | 0.02693662 |
| Lilrb3a | leukocyte immunoglobulin-like receptor, subfamily B (with TM and ITIM domains), member 3A | | | ENSRNOG00000053260 | 1.333 | 0.00021586 |
| Lilrb3l | leukocyte immunoglobulin-like receptor, subfamily B (with TM and ITIM domains), member 3-like | | | ENSRNOG00000058422 | 1.063 | 0.02738359 |
| Lilrb4 | leukocyte immunoglobulin like receptor B4 | | | ENSRNOG00000027811 | 2.081 | 4.48E-07 |
| Lilrc2 | leukocyte immunoglobulin-like receptor, subfamily C, member 2 | | | ENSRNOG00000058087 | 1.123 | 0.02885385 |
| Map2k6 | mitogen-activated protein kinase kinase 6 | | | ENSRNOG00000004437 | 0.847 | 0.01881276 |
| Ncf1 | neutrophil cytosolic factor 1 | | | ENSRNOG00000001480 | 1.746 | 8.01E-06 |
| Ncf2 | neutrophil cytosolic factor 2 | | | ENSRNOG00000028016 | 0.931 | 0.01577413 |
| Ncf4 | neutrophil cytosolic factor 4 | | | ENSRNOG00000006940 | 1.129 | 0.00118834 |
| Nfkb1 | nuclear factor kappa B subunit 1 | | | ENSRNOG00000023258 | 0.418 | 0.06528867 |
| Nfkb2 | nuclear factor kappa B subunit 2 | | | ENSRNOG00000019311 | 0.627 | 0.00144313 |
| Nfkbia | NFKB inhibitor alpha | | | ENSRNOG00000007390 | 0.338 | 0.04144616 |
| Pik3cd | phosphatidylinositol-4,5-bisphosphate 3-kinase, catalytic subunit delta | | | ENSRNOG00000016846 | 1.086 | 0.00106899 |
| Plcg2 | phospholipase C, gamma 2 | | | ENSRNOG00000051986 | 0.855 | 0.03748718 |
| Relb | RELB proto-oncogene, NF-kB subunit | | | ENSRNOG00000033235 | 1.055 | 0.00299987 |
| Sirpa | signal-regulatory protein alpha | | | ENSRNOG00000004763 | 1.223 | 0.00015422 |
| Socs3 | suppressor of cytokine signaling 3 | | | ENSRNOG00000002946 | 1.413 | 0.00438988 |
| Spi1 | Spi-1 proto-oncogene | | | ENSRNOG00000012172 | 1.239 | 0.000629 |
| Stat2 | signal transducer and activator of transcription 2 | | | ENSRNOG00000031081 | 0.716 | 2.72E-05 |
| Syk | spleen associated tyrosine kinase | | | ENSRNOG00000012160 | 0.987 | 0.00381809 |
| Tgfb1 | transforming growth factor, beta 1 | | | ENSRNOG00000020652 | 0.812 | 0.00048955 |
| Tnfrsf11a | TNF receptor superfamily member 11A | | | ENSRNOG00000015615 | 1.065 | 0.02955572 |
| Tnfrsf11b | TNF receptor superfamily member 11B | | | ENSRNOG00000008336 | 0.742 | 0.05654299 |
| Tnfrsf1a | TNF receptor superfamily member 1A | | | ENSRNOG00000031312 | 0.550 | 0.08338849 |
| Trem2 | triggering receptor expressed on myeloid cells 2 | | | ENSRNOG00000013578 | 1.143 | 0.0263419 |
| Tyrobp | Tyro protein tyrosine kinase binding protein | | | ENSRNOG00000020845 | 1.091 | 0.00258088 |
| **Rheumatoid arthritis** | | | |  |  |  |
| **Acronym** | **Gene description** | | | **ENSEMBL Reference** | **log2FC** | **Adjusted P** |
| Acp5 | acid phosphatase 5, tartrate resistant | | | ENSRNOG00000046261 | 0.476 | 0.02608932 |
| Atp6v0c | ATPase H+ transporting V0 subunit C | | | ENSRNOG00000006542 | 0.180 | 0.08549256 |
| Ccl2 | C-C motif chemokine ligand 2 | | | ENSRNOG00000007159 | 1.569 | 0.00017163 |
| Ccl3 | C-C motif chemokine ligand 3 | | | ENSRNOG00000011205 | 0.906 | 0.06197727 |
| Ccl5 | C-C motif chemokine ligand 5 | | | ENSRNOG00000010906 | 0.913 | 0.00077027 |
| Cd80 | CD80 molecule | | | ENSRNOG00000001527 | 1.409 | 0.00152716 |
| Cd86 | CD86 molecule | | | ENSRNOG00000038835 | 0.950 | 0.00354688 |
| Csf1 | colony stimulating factor 1 | | | ENSRNOG00000018659 | 0.810 | 0.02787173 |
| Ctsk | cathepsin K | | | ENSRNOG00000021155 | 0.490 | 0.01265361 |
| Icam1 | intercellular adhesion molecule 1 | | | ENSRNOG00000020679 | 1.248 | 2.44E-05 |
| Il1a | interleukin 1 alpha | | | ENSRNOG00000004575 | 1.152 | 0.0002448 |
| Il1b | interleukin 1 beta | | | ENSRNOG00000004649 | 1.714 | 2.58E-05 |
| Il18 | interleukin 18 | | | ENSRNOG00000009848 | 0.679 | 0.06437737 |
| Itgal | integrin subunit alpha L | | | ENSRNOG00000017980 | 1.504 | 0.00032999 |
| Itgb2 | integrin subunit beta 2 | | | ENSRNOG00000001224 | 0.903 | 0.01454084 |
| RT1-Ba | RT1 class II, locus Ba | | | ENSRNOG00000000451 | 1.408 | 0.00402879 |
| RT1-Bb | RT1 class II, locus Bb | | | ENSRNOG00000032708 | 1.374 | 0.00139534 |
| RT1-Da | RT1 class II, locus Da | | | ENSRNOG00000032844 | 1.307 | 0.00820569 |
| RT1-Db1 | RT1 class II, locus Db1 | | | ENSRNOG00000033215 | 1.395 | 0.00422669 |
| Tgfb1 | transforming growth factor, beta 1 | | | ENSRNOG00000020652 | 0.812 | 0.00048955 |
| Tlr2 | toll-like receptor 2 | | | ENSRNOG00000009822 | 1.650 | 2.19E-05 |
| Tnfrsf11a | TNF receptor superfamily member 11A | | | ENSRNOG00000015615 | 1.065 | 0.02955572 |
| Tnfsf13b | TNF superfamily member 13b | | | ENSRNOG00000014464 | 0.791 | 0.07036745 |
| **Systemic lupus erythematosus** | | | | |  |  |
| **Acronym** | **Gene description** | | | **ENSEMBL Reference** | **log2FC** | **Adjusted P** |
| Actn1 | actinin, alpha 1 | | | ENSRNOG00000056756 | 0.531 | 0.16004025 |
| C1qa | complement C1q A chain | | | ENSRNOG00000012807 | 0.642 | 0.06250174 |
| C1qb | complement C1q B chain | | | ENSRNOG00000012749 | 0.770 | 0.02624876 |
| C1qc | complement C1q C chain | | | ENSRNOG00000012804 | 0.725 | 0.04279283 |
| C1r | complement C1r | | | ENSRNOG00000011796 | 0.516 | 0.11635859 |
| C1s | complement C1s | | | ENSRNOG00000011971 | 0.691 | 0.0256251 |
| C3 | complement C3 | | | ENSRNOG00000046834 | 0.777 | 0.01474009 |
| C4a | complement C4A | | | ENSRNOG00000047657 | 0.121 | 0.83302637 |
| C6 | complement C6 | | | ENSRNOG00000024115 | 0.942 | 0.02349783 |
| C9 | complement C9 | | | ENSRNOG00000013736 | 1.047 | 0.01328089 |
| Cd28 | CD28 molecule | | | ENSRNOG00000010283 | 0.787 | 0.15200581 |
| Cd40 | CD40 molecule | | | ENSRNOG00000018488 | 0.916 | 0.0160465 |
| Cd80 | CD80 molecule | | | ENSRNOG00000001527 | 1.409 | 0.00152716 |
| Cd86 | CD86 molecule | | | ENSRNOG00000038835 | 0.950 | 0.00354688 |
| Fcgr1a | Fc fragment of IgG receptor Ia | | | ENSRNOG00000021199 | 1.110 | 0.00046746 |
| Fcgr3a | Fc fragment of IgG receptor IIIa | | | ENSRNOG00000024382 | 1.197 | 0.00522427 |
| Grin2a | Glutamate ionotropic receptor NMDA type subunit 2A | | | ENSRNOG00000033942 | 1.583 | 0.0012 |
| H2afv | H2A histone family, member V | | | ENSRNOG00000052275 | 0.343 | 0.00362797 |
| H2afz | H2A histone family, member Z | | | ENSRNOG00000010306 | 0.249 | 0.00852973 |
| H3f3b | H3 histone family member 3B | | | ENSRNOG00000006532 | 0.142 | 0.1464927 |
| Il10 | interleukin 10 | | | ENSRNOG00000004647 | 1.552 | 6.23E-05 |
| Mcpt8l2 | mast cell protease 8-like 2 | | | ENSRNOG00000049096 | 1.126 | 0.01042321 |
| RT1-Ba | RT1 class II, locus Ba | | | ENSRNOG00000000451 | 1.408 | 0.00402879 |
| RT1-Bb | RT1 class II, locus Bb | | | ENSRNOG00000032708 | 1.374 | 0.00139534 |
| RT1-Da | RT1 class II, locus Da | | | ENSRNOG00000032844 | 1.307 | 0.00820569 |
| RT1-Db1 | RT1 class II, locus Db1 | | | ENSRNOG00000033215 | 1.395 | 0.00422669 |
| **Asthma** |  | | |  |  |  |
| **Acronym** | **Gene description** | | | **ENSEMBL Reference** | **log2FC** | **Adjusted P** |
| Cd40 | CD40 molecule | | | ENSRNOG00000018488 | 0.916 | 0.0160465 |
| Fcer1a | Fc fragment of IgE receptor Ia | | | ENSRNOG00000009177 | 0.971 | 0.06573992 |
| Fcer1g | Fc fragment of IgE receptor Ig | | | ENSRNOG00000024159 | 1.077 | 0.00225446 |
| Il10 | interleukin 10 | | | ENSRNOG00000004647 | 1.552 | 6.23E-05 |
| Prg2 | proteoglycan 2 | | | ENSRNOG00000008394 | 1.362 | 0.00617694 |
| RT1-Ba | RT1 class II, locus Ba | | | ENSRNOG00000000451 | 1.408 | 0.00402879 |
| RT1-Bb | RT1 class II, locus Bb | | | ENSRNOG00000032708 | 1.374 | 0.00139534 |
| RT1-Da | RT1 class II, locus Da | | | ENSRNOG00000032844 | 1.307 | 0.00820569 |
| RT1-Db1 | RT1 class II, locus Db1 | | | ENSRNOG00000033215 | 1.395 | 0.00422669 |
| **Primary immunodeficiency** | | | | |  |  |
| **Acronym** | **Gene description** | | | **ENSEMBL Reference** | **log2FC** | **Adjusted P** |
| Ada | adenosine deaminase | | | ENSRNOG00000010265 | 0.520 | 0.12134469 |
| Blnk | B-cell linker | | | ENSRNOG00000013967 | 0.785 | 0.04865425 |
| Btk | Bruton tyrosine kinase | | | ENSRNOG00000052407 | 0.890 | 0.03487453 |
| Cd19 | CD19 molecule | | | ENSRNOG00000018311 | 1.180 | 0.00903096 |
| Cd3d | CD3d molecule | | | ENSRNOG00000015994 | 0.846 | 0.10305006 |
| Cd3e | CD3e molecule | | | ENSRNOG00000016069 | 0.845 | 0.04610015 |
| Cd4 | CD4 molecule | | | ENSRNOG00000016294 | 0.920 | 0.01183557 |
| Cd40 | CD40 molecule | | | ENSRNOG00000018488 | 0.916 | 0.0160465 |
| Cd79a | CD79a molecule | | | ENSRNOG00000020125 | 0.899 | 0.02174589 |
| Cd8a | CD8a molecule | | | ENSRNOG00000007178 | 0.696 | 0.07256194 |
| Icos | inducible T-cell co-stimulator | | | ENSRNOG00000046196 | 0.730 | 0.08933737 |
| Il2rg | interleukin 2 receptor subunit gamma | | | ENSRNOG00000003954 | 1.422 | 6.38E-05 |
| Il7r | interleukin 7 receptor | | | ENSRNOG00000058446 | 0.674 | 0.16913339 |
| Jak3 | Janus kinase 3 | | | ENSRNOG00000018669 | 1.462 | 7.09E-05 |
| Lck | LCK proto-oncogene, Src family tyrosine kina | | | ENSRNOG00000009705 | 0.807 | 0.00349529 |
| Ptprc | protein tyrosine phosphatase, receptor type, C | | | ENSRNOG00000000655 | 1.142 | 0.00142193 |
| Rfx5 | regulatory factor X5 | | | ENSRNOG00000021012 | 0.601 | 0.00241176 |
| Tap1 | transporter 1, ATP binding cassette subfamily B member | | | ENSRNOG00000000457 | 0.444 | 0.19276015 |
| Tnfrsf13c | TNF superfamily member 13c | | | ENSRNOG00000007664 | 0.721 | 0.20143978 |
| Ung | uracil-DNA glycosylase | | | ENSRNOG00000000692 | 0.341 | 0.00047572 |
| Zap70 | zeta chain of T cell receptor associated protein kinase 7 | | | ENSRNOG00000016995 | 0.898 | 0.01221499 |
| **Natural killer cell mediated cytotoxicity** | | | | | |  |
| **Acronym** | **Gene description** | | | **ENSEMBL Reference** | **log2FC** | **Adjusted P** |
| Cd244 | CD244 molecule | | | ENSRNOG00000004698 | 1.022 | 0.01999555 |
| Cd247 | CD247 molecule | | | ENSRNOG00000003298 | 0.646 | 0.05536135 |
| Cd48 | CD48 molecule | | | ENSRNOG00000004737 | 0.753 | 0.0093009 |
| Fas | Fas cell surface death receptor | | | ENSRNOG00000019142 | 1.155 | 0.00142758 |
| Fcer1g | Fc fragment of IgE receptor Ig | | | ENSRNOG00000024159 | 1.077 | 0.00225446 |
| Fcgr3a | Fc fragment of IgG receptor IIIa | | | ENSRNOG00000024382 | 1.197 | 0.00522427 |
| Fyn | FYN proto-oncogene, Src family tyrosine kinase | | | ENSRNOG00000000596 | 0.849 | 0.00048507 |
| Gzmb | granzyme B | | | ENSRNOG00000049976 | 1.016 | 0.02904778 |
| Hcst | hematopoietic cell signal transducer | | | ENSRNOG00000020849 | 0.878 | 0.02979094 |
| Icam1 | intercellular adhesion molecule 1 | | | ENSRNOG00000020679 | 1.248 | 2.44E-05 |
| Icam2 | intercellular adhesion molecule 2 | | | ENSRNOG00000025143 | 0.526 | 0.05967036 |
| Ifngr1 | interferon gamma receptor 1 | | | ENSRNOG00000012074 | 0.658 | 0.00243399 |
| Itgal | integrin subunit alpha L | | | ENSRNOG00000017980 | 1.504 | 0.00032999 |
| Itgb2 | integrin subunit beta 2 | | | ENSRNOG00000001224 | 0.903 | 0.01454084 |
| Klrd1 | killer cell lectin like receptor D1 | | | ENSRNOG00000060246 | 0.704 | 0.07124856 |
| Klrk1 | killer cell lectin like receptor K1 | | | ENSRNOG00000061739 | 0.932 | 0.01708985 |
| Lck | LCK proto-oncogene, Src family tyrosine kina | | | ENSRNOG00000009705 | 0.807 | 0.00349529 |
| Lcp2 | lymphocyte cytosolic protein 2 | | | ENSRNOG00000005620 | 0.798 | 0.02527559 |
| Ncr1 | natural cytotoxicity triggering receptor 1 | | | ENSRNOG00000018458 | 0.750 | 0.04856369 |
| Pik3cd | phosphatidylinositol-4,5-bisphosphate 3-kinase, catalytic subunit delta | | | ENSRNOG00000016846 | 1.086 | 0.00106899 |
| Plcg2 | phospholipase C, gamma 2 | | | ENSRNOG00000051986 | 0.855 | 0.03748718 |
| Prf1 | Perforin 1 | | | ENSRNOG00000000562 | 0.571 | 0.09305383 |
| Prkcb | protein kinase C, beta | | | ENSRNOG00000012061 | 1.040 | 0.01739716 |
| Ptk2b | protein tyrosine kinase 2 beta | | | ENSRNOG00000027839 | 1.237 | 0.0013274 |
| Ptpn6 | protein tyrosine phosphatase, non-receptor type 6 | | | ENSRNOG00000014294 | 0.475 | 0.09733397 |
| Rac2 | Rac family small GTPase 2 | | | ENSRNOG00000007350 | 1.052 | 0.00411502 |
| Sh2d1b | SH2 domain containing 1B | | | ENSRNOG00000031599 | 0.996 | 0.04685247 |
| Sh3bp2 | SH3-domain binding protein 2 | | | ENSRNOG00000013747 | 0.552 | 0.02845494 |
| Syk | spleen associated tyrosine kinase | | | ENSRNOG00000012160 | 0.987 | 0.00381809 |
| Tyrobp | Tyro protein tyrosine kinase binding protein | | | ENSRNOG00000020845 | 1.091 | 0.00258088 |
| Vav1 | vav guanine nucleotide exchange factor 1 | | | ENSRNOG00000050430 | 1.045 | 0.00414263 |
| Vav2 | vav guanine nucleotide exchange factor 2 | | | ENSRNOG00000007422 | 0.374 | 0.03411427 |
| Zap70 | zeta chain of T cell receptor associated protein kinase 70 | | | ENSRNOG00000016995 | 0.898 | 0.01221499 |
| **African trypanosomiasis** | | | |  |  |  |
| **Acronym** | **Gene description** | | | **ENSEMBL Reference** | **log2FC** | **Adjusted P** |
| Fas | Fas cell surface death receptor | | | ENSRNOG00000019142 | 1.155 | 0.00142758 |
| Hba-a2 | hemoglobin alpha, adult chain 2 | | | ENSRNOG00000047321 | 0.728 | 0.01867326 |
| Hbb | hemoglobin subunit beta | | | ENSRNOG00000058105 | 0.810 | 0.00676538 |
| Icam1 | intercellular adhesion molecule 1 | | | ENSRNOG00000020679 | 1.248 | 2.44E-05 |
| Ido2 | indoleamine 2,3-dioxygenase 2 | | | ENSRNOG00000025365 | 1.049 | 0.00422669 |
| Il10 | interleukin 10 | | | ENSRNOG00000004647 | 1.552 | 6.23E-05 |
| Il18 | interleukin 18 | | | ENSRNOG00000009848 | 0.679 | 0.06437737 |
| Il1b | interleukin 1b | | | ENSRNOG00000004649 | 1.714 | 2.58E-05 |
| Lama4 | laminin subunit alpha 4 | | | ENSRNOG00000000599 | 0.569 | 0.01006952 |
| Plcb4 | phospholipase C, beta 4 | | | ENSRNOG00000033119 | 0.437 | 0.03411427 |
| Prkcb | protein kinase C, beta | | | ENSRNOG00000012061 | 1.040 | 0.01739716 |
| Tlr9 | toll-like receptor 9 | | | ENSRNOG00000048161 | 1.331 | 0.00388139 |
| Vcam1 | vascular cell adhesion molecule 1 | | | ENSRNOG00000014333 | 0.869 | 0.00894474 |
| **Chemokine signaling** | | | |  |  |  |
| **Acronym** | **Gene description** | | | **ENSEMBL Reference** | **log2FC** | **Adjusted P** |
| Adcy4 | adenylate cyclase 4 | | | ENSRNOG00000020401 | 0.490 | 0.18038738 |
| Adcy7 | adenylate cyclase 7 | | | ENSRNOG00000014776 | 0.861 | 0.00667479 |
| Arrb2 | arrestin, beta 2 | | | ENSRNOG00000019308 | 0.768 | 0.00240746 |
| Ccl2 | C-C motif chemokine ligand 2 | | | ENSRNOG00000007159 | 1.569 | 0.00017163 |
| Ccl3 | C-C motif chemokine ligand 3 | | | ENSRNOG00000011205 | 0.906 | 0.06197727 |
| Ccl4 | C-C motif chemokine ligand 4 | | | ENSRNOG00000011406 | 0.952 | 9.00E-06 |
| Ccl5 | C-C motif chemokine ligand 5 | | | ENSRNOG00000010906 | 0.913 | 0.00077027 |
| Ccl6 | chemokine (C-C motif) ligand 6 | | | ENSRNOG00000030021 | 0.775 | 0.06144963 |
| Ccl9 | chemokine (C-C motif) ligand 9 | | | ENSRNOG00000028548 | 0.707 | 0.00213517 |
| Ccl19 | C-C motif chemokine ligand 19 | | | ENSRNOG00000015668 | 1.031 | 0.00145808 |
| Ccl21 | C-C motif chemokine ligand 21 | | | ENSRNOG00000034290 | 1.283 | 0.00191636 |
| Cx3cr1 | C-X3-C motif chemokine receptor 1 | | | ENSRNOG00000018509 | 1.103 | 0.02693662 |
| Cxcl2 | C-X-C motif chemokine ligand 2 | | | ENSRNOG00000002792 | 1.113 | 0.02823363 |
| Cxcl9 | C-X-C motif chemokine ligand 9 | | | ENSRNOG00000022242 | 0.502 | 0.07263276 |
| Cxcl10 | C-X-C motif chemokine ligand 10 | | | ENSRNOG00000022256 | 1.079 | 0.00108964 |
| Cxcl11 | C-X-C motif chemokine ligand 11 | | | ENSRNOG00000022298 | 0.404 | 0.1446508 |
| Cxcl13 | C-X-C motif chemokine ligand 13 | | | ENSRNOG00000024899 | 1.407 | 0.0043727 |
| Cxcl14 | C-X-C motif chemokine ligand 14 | | | ENSRNOG00000011984 | 0.678 | 0.00765963 |
| Cxcl16 | C-X-C motif chemokine ligand 16 | | | ENSRNOG00000026647 | 1.404 | 1.93E-05 |
| Cxcr2 | C-X-C motif chemokine receptor 2 | | | ENSRNOG00000014269 | 1.329 | 0.00796676 |
| Cxcr4 | C-X-C motif chemokine receptor 4 | | | ENSRNOG00000003866 | 1.319 | 6.87E-05 |
| Dock2 | dedicator of cytokinesis 2 | | | ENSRNOG00000006932 | 0.681 | 0.15425969 |
| Elmo1 | engulfment and cell motility 1 | | | ENSRNOG00000059705 | 0.826 | 0.03746427 |
| Fgr | FGR proto-oncogene, Src family tyrosine kinase | | | ENSRNOG00000009912 | 1.353 | 0.00035326 |
| Gnai1 | G protein subunit alpha i1 | | | ENSRNOG00000057096 | 0.523 | 0.0270346 |
| Gnai2 | G protein subunit alpha i2 | | | ENSRNOG00000016592 | 0.522 | 0.01055457 |
| Gnb2 | G protein subunit beta 2 | | | ENSRNOG00000001409 | 0.246 | 0.06250174 |
| Gnb4 | G protein subunit beta 4 | | | ENSRNOG00000011070 | 0.654 | 0.14397774 |
| Gng2 | G protein subunit gamma 2 | | | ENSRNOG00000048980 | 0.825 | 0.01929356 |
| Gngt2 | G protein subunit gamma transducin 2 | | | ENSRNOG00000006108 | 1.057 | 0.00281874 |
| Grk3 | G protein-coupled receptor kinase 3 | | | ENSRNOG00000059456 | 1.020 | 0.01504335 |
| Hck | HCK proto-oncogene, Src family tyrosine kinase | | | ENSRNOG00000009331 | 1.128 | 0.00496442 |
| Ikbkb | inhibitor of nuclear factor kappa B kinase subunit beta | | | ENSRNOG00000019073 | 0.374 | 0.08821073 |
| Itk | IL2-inducible T-cell kinase | | | ENSRNOG00000006860 | 0.763 | 0.03536247 |
| Jak2 | Janus kinase 2 | | | ENSRNOG00000059968 | 0.564 | 0.08130277 |
| Jak3 | Janus kinase 3 | | | ENSRNOG00000018669 | 1.462 | 7.09E-05 |
| Lyn | LYN proto-oncogene, Src family tyrosine kinase | | | ENSRNOG00000008180 | 0.681 | 0.01897376 |
| Ncf1 | neutrophil cytosolic factor 1 | | | ENSRNOG00000001480 | 1.746 | 8.01E-06 |
| Nfkb1 | nuclear factor kappa B subunit 1 | | | ENSRNOG00000023258 | 0.418 | 0.06528867 |
| Nfkbia | NFKB inhibitor alpha | | | ENSRNOG00000007390 | 0.338 | 0.04144616 |
| Nfkbib | NFKB inhibitor beta | | | ENSRNOG00000020063 | 0.243 | 0.10595847 |
| Pf4 | platelet factor 4 | | | ENSRNOG00000028015 | 0.678 | 0.05159754 |
| Pik3cd | phosphatidylinositol-4,5-bisphosphate 3-kinase, catalytic subunit delta | | | ENSRNOG00000016846 | 1.086 | 0.00106899 |
| Pik3cg | phosphatidylinositol-4,5-bisphosphate 3-kinase, catalytic subunit gamma | | | ENSRNOG00000009385 | 0.770 | 0.02536242 |
| Pik3r5 | phosphoinositide-3-kinase, regulatory subunit 5 | | | ENSRNOG00000023428 | 1.132 | 0.00180089 |
| Pik3r6 | phosphoinositide-3-kinase, regulatory subunit 6 | | | ENSRNOG00000003992 | 0.456 | 0.05116291 |
| Plcb2 | phospholipase C, beta 2 | | | ENSRNOG00000058337 | 0.509 | 0.17140923 |
| Plcb4 | phospholipase C, beta 4 | | | ENSRNOG00000033119 | 0.437 | 0.03411427 |
| Prex1 | phosphatidylinositol-3,4,5-trisphosphate-dependent Rac exchange factor 1 | | | ENSRNOG00000006952 | 1.090 | 0.00573769 |
| Prkcb | protein kinase C, beta | | | ENSRNOG00000012061 | 1.040 | 0.01739716 |
| Prkcd | protein kinase C, delta | | | ENSRNOG00000016346 | 0.494 | 0.00381496 |
| Prkcz | protein kinase C, zeta | | | ENSRNOG00000015480 | 0.572 | 0.00351668 |
| Ptk2b | protein tyrosine kinase 2 beta | | | ENSRNOG00000027839 | 1.237 | 0.0013274 |
| Rac2 | Rac family small GTPase 2 | | | ENSRNOG00000007350 | 1.052 | 0.00411502 |
| Rasgrp2 | RAS guanyl releasing protein 2 | | | ENSRNOG00000021098 | 0.249 | 0.14006126 |
| Src | SRC proto-oncogene, non-receptor tyrosine kinase | | | ENSRNOG00000009495 | 0.877 | 0.00544661 |
| Stat1 | signal transducer and activator of transcription 1 | | | ENSRNOG00000014079 | 0.492 | 0.16315358 |
| Stat2 | signal transducer and activator of transcription 2 | | | ENSRNOG00000031081 | 0.716 | 2.72E-05 |
| Vav1 | vav guanine nucleotide exchange factor 1 | | | ENSRNOG00000050430 | 1.045 | 0.00414263 |
| Vav2 | vav guanine nucleotide exchange factor 2 | | | ENSRNOG00000007422 | 0.374 | 0.03411427 |
| Vav3 | vav guanine nucleotide exchange factor 3 | | | ENSRNOG00000020485 | 0.429 | 0.12420858 |
| Was | WASP actin nucleation promoting factor | | | ENSRNOG00000031058 | 0.762 | 0.03368191 |
| Xcl1 | X-C motif chemokine ligand 1 | | | ENSRNOG00000002964 | 0.585 | 0.11036923 |
| **Cytosolic DNA-sensing** | | | |  |  |  |
| **Acronym** | **Gene description** | | | **ENSEMBL Reference** | **log2FC** | **Adjusted P** |
| Aim2 | absent in melanoma 2 | | | ENSRNOG00000003480 | 0.813 | 0.03154279 |
| Casp1 | caspase 1 | | | ENSRNOG00000007372 | 0.683 | 0.10786101 |
| Ccl4 | C-C motif chemokine ligand 4 | | | ENSRNOG00000011406 | 0.952 | 9.00E-06 |
| Ccl5 | C-C motif chemokine ligand 5 | | | ENSRNOG00000010906 | 0.913 | 0.00077027 |
| Cxcl10 | C-X-C motif chemokine ligand 10 | | | ENSRNOG00000022256 | 1.079 | 0.00108964 |
| Ikbkb | inhibitor of nuclear factor kappa B kinase subunit beta | | | ENSRNOG00000019073 | 0.374 | 0.08821073 |
| Ikbke | inhibitor of nuclear factor kappa B kinase subunit epsilon | | | ENSRNOG00000025100 | 1.058 | 0.00093628 |
| Il18 | interleukin 18 | | | ENSRNOG00000009848 | 0.679 | 0.06437737 |
| Il1b | interleukin 1 beta | | | ENSRNOG00000004649 | 1.714 | 2.58E-05 |
| Il33 | interleukin 33 | | | ENSRNOG00000016456 | 1.645 | 1.71E-07 |
| Nfkb1 | nuclear factor kappa B subunit 1 | | | ENSRNOG00000023258 | 0.418 | 0.06528867 |
| Nfkbia | NFKB inhibitor alpha | | | ENSRNOG00000007390 | 0.338 | 0.04144616 |
| Nfkbib | NFKB inhibitor beta | | | ENSRNOG00000020063 | 0.243 | 0.10595847 |
| Polr3h | RNA polymerase III subunit H | | | ENSRNOG00000004471 | 0.313 | 0.07851258 |
| Pycard | PYD and CARD domain containing | | | ENSRNOG00000019675 | 1.027 | 0.01196718 |
| Ripk3 | receptor-interacting serine-threonine kinase 3 | | | ENSRNOG00000020465 | 0.924 | 0.03839795 |
| Tmem173 | transmembrane protein 173 | | | ENSRNOG00000042137 | 1.417 | 4.96E-05 |
| Trex1 | three prime repair exonuclease 1 | | | ENSRNOG00000022540 | 0.420 | 0.01724867 |
| Zbp1 | Z-DNA binding protein 1 | | | ENSRNOG00000006314 | 0.887 | 0.03035486 |
| **Intestinal immune network for IgA production** | | | | | |  |
| **Acronym** | **Gene description** | | | **ENSEMBL Reference** | **log2FC** | **Adjusted P** |
| Cd28 | CD28 molecule | | | ENSRNOG00000010283 | 0.787 | 0.15200581 |
| Cd40 | CD40 molecule | | | ENSRNOG00000018488 | 0.916 | 0.0160465 |
| Cd80 | CD80 molecule | | | ENSRNOG00000001527 | 1.409 | 0.00152716 |
| Cd86 | CD86 molecule | | | ENSRNOG00000038835 | 0.950 | 0.00354688 |
| Cxcr4 | C-X-C motif chemokine receptor 4 | | | ENSRNOG00000003866 | 1.319 | 6.87E-05 |
| Icos | inducible T-cell co-stimulator | | | ENSRNOG00000046196 | 0.730 | 0.08933737 |
| Il10 | interleukin 10 | | | ENSRNOG00000004647 | 1.552 | 6.23E-05 |
| Il15ra | Interleukin 15 receptor subunit alpha | | | ENSRNOG00000018706 | 0.561 | 6.07E-05 |
| Itga4 | integrin subunit alpha 4 | | | ENSRNOG00000004861 | 0.780 | 0.08536026 |
| Itgb7 | integrin subunit beta 7 | | | ENSRNOG00000012208 | 0.900 | 0.01428388 |
| Ltbr | Lymphotoxin beta receptor | | | ENSRNOG00000019264 | 0.284 | 0.02979056 |
| RT1-Ba | RT1 class II, locus Ba | | | ENSRNOG00000000451 | 1.408 | 0.00402879 |
| RT1-Bb | RT1 class II, locus Bb | | | ENSRNOG00000032708 | 1.374 | 0.00139534 |
| RT1-Da | RT1 class II, locus Da | | | ENSRNOG00000032844 | 1.307 | 0.00820569 |
| RT1-Db1 | RT1 class II, locus Db1 | | | ENSRNOG00000033215 | 1.395 | 0.00422669 |
| Tgfb1 | transforming growth factor, beta 1 | | | ENSRNOG00000020652 | 0.812 | 0.00048955 |
| Tnfrsf17 | TNF superfamily member 17 | | | ENSRNOG00000021987 | 0.815 | 0.1372405 |
| Tnfsf13b | TNF superfamily member 13b | | | ENSRNOG00000014464 | 0.791 | 0.07036745 |
| **Tuberculosis** | | | |  |  |  |
| **Acronym** | **Gene description** | | | **ENSEMBL Reference** | **log2FC** | **Adjusted P** |
| Atp6v0c | ATPase H+ transporting V0 subunit C | | | ENSRNOG00000006542 | 0.180 | 0.08549256 |
| Bcl10 | BCL10, immune signaling adaptor | | | ENSRNOG00000042389 | 0.342 | 0.02624876 |
| C3 | complement C3 | | | ENSRNOG00000046834 | 0.777 | 0.01474009 |
| Camk2b | calcium/calmodulin-dependent protein kinase II beta | | | ENSRNOG00000052080 | 1.373 | 0.00588558 |
| Camk2d | calcium/calmodulin-dependent protein kinase II delta | | | ENSRNOG00000011589 | 0.703 | 0.05185511 |
| Camp | cathelicidin antimicrobial peptide | | | ENSRNOG00000020733 | 0.887 | 0.07959316 |
| Cd74 | CD74 molecule | | | ENSRNOG00000018735 | 1.655 | 4.74E-05 |
| Clec4e | C-type lectin domain family 4, member E | | | ENSRNOG00000061394 | 1.895 | 5.87E-05 |
| Clec7a | C-type lectin domain containing 7A | | | ENSRNOG00000054251 | 1.954 | 4.47E-08 |
| Coro1a | coronin 1A | | | ENSRNOG00000019430 | 1.217 | 3.98E-06 |
| Ctss | Cathepsin S | | | ENSRNOG00000021157 | 0.803 | 0.04312882 |
| Fcer1g | Fc fragment of IgG receptor Ig | | | ENSRNOG00000024159 | 1.077 | 0.00225446 |
| Fcgr3a | Fc fragment of IgG receptor IIIa | | | ENSRNOG00000024382 | 1.197 | 0.00522427 |
| Ifngr1 | interferon gamma receptor 1 | | | ENSRNOG00000012074 | 0.658 | 0.00243399 |
| Il10 | interleukin 10 | | | ENSRNOG00000004647 | 1.552 | 6.23E-05 |
| Il10ra | interleukin 10 receptor subunit alpha | | | ENSRNOG00000016308 | 1.107 | 0.00427164 |
| Il10rb | interleukin 10 receptor subunit beta | | | ENSRNOG00000028638 | 0.453 | 0.01243751 |
| Il1a | interleukin 1 alpha | | | ENSRNOG00000004575 | 1.152 | 0.0002448 |
| Il1b | interleukin 1 beta | | | ENSRNOG00000004649 | 1.714 | 2.58E-05 |
| Itgax | integrin subunit alpha X | | | ENSRNOG00000036703 | 1.078 | 1.69E-05 |
| Jak2 | Janus kinase 2 | | | ENSRNOG00000059968 | 0.564 | 0.08130277 |
| Lamp1 | lysosomal-associated membrane protein 1 | | | ENSRNOG00000019629 | 0.156 | 0.05284966 |
| Lbp | lipopolysaccharide binding protein | | | ENSRNOG00000014532 | 1.265 | 0.01227787 |
| Lsp1 | lymphocyte-specific protein 1 | | | ENSRNOG00000020300 | 0.937 | 5.45E-08 |
| Mrc2 | mannose receptor, C type 2 | | | ENSRNOG00000006548 | 0.953 | 0.00713081 |
| Nfkb1 | nuclear factor kappa B subunit 1 | | | ENSRNOG00000023258 | 0.418 | 0.06528867 |
| Nfyb | nuclear transcription factor Y subunit beta | | | ENSRNOG00000010309 | 0.407 | 0.0037893 |
| Nod2 | nucleotide-binding oligomerization domain containing 2 | | | ENSRNOG00000014124 | 1.248 | 0.0033119 |
| Rfx5 | regulatory factor X5 | | | ENSRNOG00000021012 | 0.601 | 0.00241176 |
| Ripk2 | receptor-interacting serine-threonine kinase 2 | | | ENSRNOG00000009389 | 0.505 | 0.08523801 |
| RT1-Ba | RT1 class II, locus Ba | | | ENSRNOG00000000451 | 1.408 | 0.00402879 |
| RT1-Bb | RT1 class II, locus Bb | | | ENSRNOG00000032708 | 1.374 | 0.00139534 |
| RT1-Da | RT1 class II, locus Da | | | ENSRNOG00000032844 | 1.307 | 0.00820569 |
| RT1-Db1 | RT1 class II, locus Db1 | | | ENSRNOG00000033215 | 1.395 | 0.00422669 |
| Src | SRC proto-oncogene, non-receptor tyrosine kinase | | | ENSRNOG00000009495 | 0.877 | 0.00544661 |
| Syk | spleen associated tyrosine kinase | | | ENSRNOG00000012160 | 0.987 | 0.00381809 |
| Tgfb1 | transforming growth factor, beta 1 | | | ENSRNOG00000020652 | 0.812 | 0.00048955 |
| Tlr2 | toll-like receptor 2 | | | ENSRNOG00000009822 | 1.650 | 2.19E-05 |
| Tlr9 | toll-like receptor 9 | | | ENSRNOG00000048161 | 1.331 | 0.00388139 |
| Tnfrsf1a | TNF receptor superfamily member 1A | | | ENSRNOG00000031312 | 0.550 | 0.08338849 |
| **Pertussis** |  | | |  |  |  |
| **Acronym** | **Gene description** | | | **ENSEMBL Reference** | **log2FC** | **Adjusted P** |
| C1qa | complement C1q A chain | | | ENSRNOG00000012807 | 0.642 | 0.06250174 |
| C1qb | complement C1q B chain | | | ENSRNOG00000012749 | 0.770 | 0.02624876 |
| C1qc | complement C1q C chain | | | ENSRNOG00000012804 | 0.725 | 0.04279283 |
| C1r | complement C1r | | | ENSRNOG00000011796 | 0.516 | 0.11635859 |
| C1s | complement C1s | | | ENSRNOG00000011971 | 0.691 | 0.0256251 |
| C2 | complement C2 | | | ENSRNOG00000051235 | 0.288 | 0.37851584 |
| C3 | complement C3 | | | ENSRNOG00000046834 | 0.777 | 0.01474009 |
| C4a | complement C4A | | | ENSRNOG00000047657 | 0.121 | 0.83302637 |
| C4bpa | complement component 4 binding protein, alpha | | | ENSRNOG00000004062 | 0.821 | 0.00996253 |
| C4bpb | complement component 4 binding protein, beta | | | ENSRNOG00000004125 | 0.934 | 4.78E-05 |
| C5 | complement C5 | | | ENSRNOG00000018899 | 0.305 | 0.3515497 |
| Calm3 | calmodulin 3 | | | ENSRNOG00000004060 | 0.227 | 0.28763901 |
| Casp3 | caspase 3 | | | ENSRNOG00000010475 | 0.252 | 0.30184544 |
| Casp7 | caspase 7 | | | ENSRNOG00000056216 | 0.321 | 0.00020047 |
| Cfl1 | cofilin 1 | | | ENSRNOG00000020660 | 0.307 | 0.08333429 |
| Fos | Fos proto-oncogene, AP-1 transcription factor subunit | | | ENSRNOG00000008015 | 0.674 | 0.15874707 |
| Gnai1 | G protein subunit alpha i1 | | | ENSRNOG00000057096 | 0.523 | 0.0270346 |
| Gnai2 | G protein subunit alpha i2 | | | ENSRNOG00000016592 | 0.522 | 0.01055457 |
| Il10 | interleukin 10 | | | ENSRNOG00000004647 | 1.552 | 6.23E-05 |
| Il1a | interleukin 1 alpha | | | ENSRNOG00000004575 | 1.152 | 0.0002448 |
| Il1b | interleukin 1 beta | | | ENSRNOG00000004649 | 1.714 | 2.58E-05 |
| Irak4 | interleukin-1 receptor-associated kinase 4 | | | ENSRNOG00000005965 | 0.315 | 0.35937686 |
| Irf1 | interferon regulatory factor 1 | | | ENSRNOG00000008144 | 0.536 | 0.29159302 |
| Irf8 | interferon regulatory factor 8 | | | ENSRNOG00000017869 | 0.807 | 0.00759497 |
| Itgb2 | integrin subunit beta 2 | | | ENSRNOG00000001224 | 0.903 | 0.01454084 |
| Jun | Jun proto-oncogene, AP-1 transcription factor subunit | | | ENSRNOG00000026293 | 0.413 | 0.3852847 |
| Mapk11 | mitogen-activated protein kinase 11 | | | ENSRNOG00000006984 | 0.332 | 0.27657293 |
| Myd88 | MYD88, innate immune signal transduction adaptor | | | ENSRNOG00000013634 | 0.175 | 0.2955869 |
| Nfkb1 | nuclear factor kappa B subunit 1 | | | ENSRNOG00000023258 | 0.418 | 0.06528867 |
| Nlrp3 | NLR family, pyrin domain containing 3 | | | ENSRNOG00000003170 | 0.778 | 0.0525725 |
| Nod1 | nucleotide-binding oligomerization domain containing 1 | | | ENSRNOG00000010629 | 0.377 | 0.14080154 |
| Serping1 | serpin family G member 1 | | | ENSRNOG00000007457 | 0.601 | 0.06757624 |
| Ticam2 | toll-like receptor adaptor molecule 2 | | | ENSRNOG00000042070 | 0.478 | 0.3116609 |
| Tlr4 | toll-like receptor 4 | | | ENSRNOG00000010522 | 0.549 | 0.27371904 |
| **Chagas disease (American trypanosomiasis)** | | | | | |  |
| **Acronym** | **Gene description** | | | **ENSEMBL Reference** | **log2FC** | **Adjusted P** |
| Ace3 | angiotensin I converting enzyme (peptidyl-dipeptidase A) 3 | | | ENSRNOG00000007467 | 1.069 | 0.00345673 |
| Bdkrb2 | bradykinin receptor B2 | | | ENSRNOG00000047300 | 0.846 | 0.12133858 |
| C1qa | complement C1q A chain | | | ENSRNOG00000012807 | 0.642 | 0.06250174 |
| C1qb | complement C1q B chain | | | ENSRNOG00000012749 | 0.770 | 0.02624876 |
| C1qc | complement C1q C chain | | | ENSRNOG00000012804 | 0.725 | 0.04279283 |
| C3 | complement C3 | | | ENSRNOG00000046834 | 0.777 | 0.01474009 |
| Ccl2 | C-C motif chemokine ligand 2 | | | ENSRNOG00000007159 | 1.569 | 0.00017163 |
| Ccl3 | C-C motif chemokine ligand 3 | | | ENSRNOG00000011205 | 0.906 | 0.06197727 |
| Ccl5 | C-C motif chemokine ligand 5 | | | ENSRNOG00000010906 | 0.913 | 0.00077027 |
| Cd247 | CD247 molecule | | | ENSRNOG00000003298 | 0.646 | 0.05536135 |
| Cd3d | CD3d molecule | | | ENSRNOG00000015994 | 0.846 | 0.10305006 |
| Cd3e | CD3e molecule | | | ENSRNOG00000016069 | 0.845 | 0.04610015 |
| Cd3g | CD3g molecule | | | ENSRNOG00000015945 | 0.907 | 0.06376473 |
| Fas | Fas cell surface death receptor | | | ENSRNOG00000019142 | 1.155 | 0.00142758 |
| Gna14 | G protein subunit alpha 14 | | | ENSRNOG00000014840 | 0.716 | 0.0950869 |
| Gna15 | G protein subunit alpha 15 | | | ENSRNOG00000005378 | 1.318 | 8.42E-05 |
| Gnai1 | G protein subunit alpha i1 | | | ENSRNOG00000057096 | 0.523 | 0.0270346 |
| Gnai2 | G protein subunit alpha i2 | | | ENSRNOG00000016592 | 0.522 | 0.01055457 |
| Gnal | G protein subunit alpha L | | | ENSRNOG00000010440 | 0.844 | 0.12204556 |
| Ifngr1 | interferon gamma receptor 1 | | | ENSRNOG00000012074 | 0.658 | 0.00243399 |
| Ikbkb | inhibitor of nuclear factor kappa B kinase subunit beta | | | ENSRNOG00000019073 | 0.374 | 0.08821073 |
| Il10 | interleukin 10 | | | ENSRNOG00000004647 | 1.552 | 6.23E-05 |
| Il1b | interleukin 1 beta | | | ENSRNOG00000004649 | 1.714 | 2.58E-05 |
| Nfkb1 | nuclear factor kappa B subunit 1 | | | ENSRNOG00000023258 | 0.418 | 0.06528867 |
| Nfkbia | NFKB inhibitor alpha | | | ENSRNOG00000007390 | 0.338 | 0.04144616 |
| Pik3cd | phosphatidylinositol-4,5-bisphosphate 3-kinase, catalytic subunit delta | | | ENSRNOG00000016846 | 1.086 | 0.00106899 |
| Plcb4 | phospholipase C, beta 4 | | | ENSRNOG00000033119 | 0.437 | 0.03411427 |
| Serpine1 | serpin family G member 1 | | | ENSRNOG00000001414 | 1.898 | 1.20E-18 |
| Tgfb1 | transforming growth factor, beta 1 | | | ENSRNOG00000020652 | 0.812 | 0.00048955 |
| Tlr2 | toll-like receptor 2 | | | ENSRNOG00000009822 | 1.650 | 2.19E-05 |
| Tlr9 | toll-like receptor 9 | | | ENSRNOG00000048161 | 1.331 | 0.00388139 |
| Tnfrsf1a | Tnfrsf1a | | | ENSRNOG00000031312 | 0.550 | 0.08338849 |
| **C-type lectin receptor signaling** | | | | |  |  |
| **Acronym** | **Gene description** | | | **ENSEMBL Reference** | **log2FC** | **Adjusted P** |
| Bcl10 | BCL10, immune signaling adaptor | | | ENSRNOG00000042389 | 0.342 | 0.02624876 |
| Bcl3 | BCL3, transcription coactivat | | | ENSRNOG00000043416 | 1.296 | 0.00169758 |
| Card9 | caspase recruitment domain family, member 9 | | | ENSRNOG00000051470 | 0.696 | 0.11718056 |
| Casp1 | caspase 1 | | | ENSRNOG00000007372 | 0.683 | 0.10786101 |
| Clec4d | C-type lectin domain family 4, member D | | | ENSRNOG00000010181 | 1.327 | 0.00542184 |
| Clec4e | C-type lectin domain family 4, member E | | | ENSRNOG00000061394 | 1.895 | 5.87E-05 |
| Clec7a | C-type lectin domain containing 7A | | | ENSRNOG00000054251 | 1.954 | 4.47E-08 |
| Egr2 | early growth response 2 | | | ENSRNOG00000000640 | 0.850 | 0.0902866 |
| Egr3 | early growth response 3 | | | ENSRNOG00000017828 | 1.213 | 0.00759434 |
| Fcer1g | Fc fragment of IgG receptor Ig | | | ENSRNOG00000024159 | 1.077 | 0.00225446 |
| Ikbkb | inhibitor of nuclear factor kappa B kinase subunit beta | | | ENSRNOG00000019073 | 0.374 | 0.08821073 |
| Ikbke | inhibitor of nuclear factor kappa B kinase subunit epsilon | | | ENSRNOG00000025100 | 1.058 | 0.00093628 |
| Il10 | interleukin 10 | | | ENSRNOG00000004647 | 1.552 | 6.23E-05 |
| Il1b | interleukin 1 beta | | | ENSRNOG00000004649 | 1.714 | 2.58E-05 |
| Itpr3 | inositol 1,4,5-trisphosphate receptor, type 3 | | | ENSRNOG00000052795 | 0.600 | 0.04179986 |
| Lsp1 | lymphocyte-specific protein 1 | | | ENSRNOG00000020300 | 0.937 | 5.45E-08 |
| Mras | muscle RAS oncogene homolog | | | ENSRNOG00000014060 | 0.533 | 0.0272179 |
| Nfkb1 | nuclear factor kappa B subunit 1 | | | ENSRNOG00000023258 | 0.418 | 0.06528867 |
| Nfkb2 | nuclear factor kappa B subunit 2 | | | ENSRNOG00000019311 | 0.627 | 0.00144313 |
| Nfkbia | NFKB inhibitor alpha | | | ENSRNOG00000007390 | 0.338 | 0.04144616 |
| Nlrp3 | NLR family, pyrin domain containing 3 | | | ENSRNOG00000003170 | 0.778 | 0.0525725 |
| Pik3cd | phosphatidylinositol-4,5-bisphosphate 3-kinase, catalytic subunit delta | | | ENSRNOG00000016846 | 1.086 | 0.00106899 |
| Plcg2 | phospholipase C, gamma 2 | | | ENSRNOG00000051986 | 0.855 | 0.03748718 |
| Prkcd | protein kinase C, delta | | | ENSRNOG00000016346 | 0.494 | 0.00381496 |
| Pycard | PYD and CARD domain containing | | | ENSRNOG00000019675 | 1.027 | 0.01196718 |
| Relb | RELB proto-oncogene, NF-kB subunit | | | ENSRNOG00000033235 | 1.055 | 0.00299987 |
| Rras | RAS related | | | ENSRNOG00000037247 | 0.525 | 0.0007021 |
| Rras2 | RAS related 2 | | | ENSRNOG00000012258 | 0.407 | 0.1013708 |
| Src | SRC proto-oncogene, non-receptor tyrosine kinase | | | ENSRNOG00000009495 | 0.877 | 0.00544661 |
| Stat2 | signal transducer and activator of transcription 2 | | | ENSRNOG00000031081 | 0.716 | 2.72E-05 |
| Syk | spleen associated tyrosine kinase | | | ENSRNOG00000012160 | 0.987 | 0.00381809 |
| **Maturity onset diabetes of the young** | | | | |  |  |
| **Acronym** | **Gene description** | | | **ENSEMBL Reference** | **log2FC** | **Adjusted P** |
| Gck | glucokinase | | | ENSRNOG00000061527 | -2.460 | 3.37E-08 |
| Hes1 | hes family bHLH transcription factor 1 | | | ENSRNOG00000001720 | -0.912 | 0.00121684 |
| Hnf1a | HNF1 homeobox A | | | ENSRNOG00000001183 | -0.450 | 0.00196918 |
| Hnf4a | hepatocyte nuclear factor 4, alpha | | | ENSRNOG00000008895 | -0.366 | 0.01170365 |
| Hnf4g | hepatocyte nuclear factor 4, gamma | | | ENSRNOG00000008971 | -1.846 | 1.05E-16 |
| **Leukocyte transendothelial migration** | | | | |  |  |
| **Acronym** | **Gene description** | | | **ENSEMBL Reference** | **log2FC** | **Adjusted P** |
| Afdn | afadin, adherens junction formation factor | | | ENSRNOG00000023753 | -0.422 | 0.00036817 |
| Arhgap5 | Rho GTPase activating protein 5 | | | ENSRNOG00000004696 | -0.682 | 4.81E-06 |
| Cldn1 | claudin 1 | | | ENSRNOG00000001926 | -3.148 | 4.35E-172 |
| Cldn2 | claudin 2 | | | ENSRNOG00000054495 | -0.588 | 9.57E-06 |
| Cldn14 | claudin 14 | | | ENSRNOG00000001691 | -0.783 | 0.00039772 |
| Pik3cb | phosphatidylinositol-4,5-bisphosphate 3-kinase, catalytic subunit beta | | | ENSRNOG00000016384 | -0.547 | 0.00016849 |
| Rapgef4 | Rap guanine nucleotide exchange factor 4 | | | ENSRNOG00000001516 | -0.740 | 0.00108122 |
| Vcl | vinculin | | | ENSRNOG00000010765 | -0.614 | 1.11E-08 |
| **Aminoacyl-tRNA biosynthesis** | | | | |  |  |
| **Acronym** | **Gene description** | | | **ENSEMBL Reference** | **log2FC** | **Adjusted P** |
| Aars2 | alanyl-tRNA synthetase 2, mitochondrial | | | ENSRNOG00000025808 | -0.172 | 0.2506597 |
| Cars | cysteinyl-tRNA synthetase | | | ENSRNOG00000020651 | -0.504 | 6.99E-05 |
| Cars2 | cysteinyl-tRNA synthetase 2, mitochondrial | | | ENSRNOG00000014526 | -0.207 | 0.32597139 |
| Dars | aspartyl-tRNA synthetase | | | ENSRNOG00000003743 | -0.246 | 0.03220231 |
| Ears2 | glutamyl-tRNA synthetase 2, mitochondrial | | | ENSRNOG00000025353 | -0.801 | 7.72E-07 |
| Farsa | phenylalanyl-tRNA synthetase subunit alpha | | | ENSRNOG00000003149 | -0.186 | 0.20442684 |
| Farsb | phenylalanyl-tRNA synthetase subunit beta | | | ENSRNOG00000014119 | -0.241 | 0.20242122 |
| Gars | glycyl-tRNA synthetase | | | ENSRNOG00000011052 | -0.380 | 0.00055007 |
| Gatb | glutamyl-tRNA amidotransferase subunit B | | | ENSRNOG00000037655 | -0.297 | 0.01447805 |
| Gatc | glutamyl-tRNA amidotransferase subunit C | | | ENSRNOG00000001161 | -0.163 | 0.31941513 |
| Hars2 | histidyl-tRNA synthetase 2, mitochondrial | | | ENSRNOG00000016087 | -0.138 | 0.30606035 |
| Iars | isoleucyl-tRNA synthetase 1 | | | ENSRNOG00000014616 | -0.485 | 9.30E-05 |
| Iars2 | isoleucyl-tRNA synthetase 2 | | | ENSRNOG00000002368 | -0.228 | 0.01503059 |
| Lars | leucyl-tRNA synthetase 1 | | | ENSRNOG00000018304 | -0.254 | 0.02830298 |
| Lars2 | leucyl-tRNA synthetase 2, mitochondrial | | | ENSRNOG00000004760 | -0.203 | 0.28344954 |
| Mars | methionyl-tRNA synthetase 1 | | | ENSRNOG00000025459 | -0.327 | 0.01451398 |
| Mtfmt | mitochondrial methionyl-tRNA formyltransferase | | | ENSRNOG00000014602 | -0.222 | 0.1002611 |
| Nars | asparaginyl-tRNA synthetase 1 | | | ENSRNOG00000017852 | -0.319 | 0.00628567 |
| Nars2 | asparaginyl-tRNA synthetase 2, mitochondrial | | | ENSRNOG00000011476 | -0.408 | 0.00297705 |
| Sars | seryl-tRNA synthetase 1 | | | ENSRNOG00000020255 | -0.223 | 0.07687219 |
| Tarsl2 | threonyl-tRNA synthetase 3 | | | ENSRNOG00000024460 | -0.843 | 0.00014428 |
| Vars | valyl-tRNA synthetase 1 | | | ENSRNOG00000000867 | -0.193 | 0.07284732 |
| Vars2 | valyl-tRNA synthetase 2, mitochondrial | | | ENSRNOG00000000833 | -0.157 | 0.33081734 |
| Wars2 | tryptophanyl tRNA synthetase 2 (mitochondrial) | | | ENSRNOG00000019508 | -0.408 | 0.01668633 |
| Yars2 | tyrosyl-tRNA synthetase 2 | | | ENSRNOG00000025252 | -0.166 | 0.27357929 |
| **Lysine degradation** | | | |  |  |  |
| **Acronym** | **Gene description** | | | **ENSEMBL Reference** | **log2FC** | **Adjusted P** |
| Acat1 | acetyl-CoA acetyltransferase 1 | | | ENSRNOG00000007862 | -0.651 | 0.03091885 |
| Aldh2 | aldehyde dehydrogenase 2 family member | | | ENSRNOG00000001344 | -0.297 | 0.0243679 |
| Bbox1 | gamma-butyrobetaine hydroxylase 1 | | | ENSRNOG00000059519 | -0.680 | 0.00621703 |
| Dot1l | DOT1 like histone lysine methyltransferase | | | ENSRNOG00000032546 | -0.660 | 3.56E-06 |
| Ehhadh | Enoyl-CoA hydratase and 3-hydroxyacyl CoA dehydrogenase | | | ENSRNOG00000001770 | -1.317 | 4.61E-05 |
| Ehmt1 | euchromatic histone lysine methyltransferase 1 | | | ENSRNOG00000007242 | -0.345 | 0.075973 |
| Hadh | hydroxyacyl-CoA dehydrogenase | | | ENSRNOG00000010697 | -0.586 | 0.04364221 |
| Hadha | hydroxyacyl-CoA dehydrogenase trifunctional multienzyme complex subunit alpha | | | ENSRNOG00000024629 | -0.467 | 0.00285055 |
| Kmt2a | lysine methyltransferase 2A | | | ENSRNOG00000015133 | -0.498 | 0.00412253 |
| Kmt2c | lysine methyltransferase 2C | | | ENSRNOG00000061080 | -0.473 | 0.06876866 |
| Kmt2d | lysine methyltransferase 2D | | | ENSRNOG00000061499 | -0.347 | 1.40E-05 |
| Kmt5a | lysine methyltransferase 5A | | | ENSRNOG00000001062 | -0.297 | 0.14904782 |
| Kmt5b | lysine methyltransferase 5B | | | ENSRNOG00000016790 | -0.234 | 0.01532953 |
| Nsd1 | nuclear receptor binding SET domain protein 1 | | | ENSRNOG00000016680 | -0.351 | 0.03328317 |
| Nsd2 | nuclear receptor binding SET domain protein 2 | | | ENSRNOG00000038140 | -0.410 | 0.00664173 |
| Nsd3 | nuclear receptor binding SET domain protein 3 | | | ENSRNOG00000015621 | -0.256 | 0.12276864 |
| Pipox | pipecolic acid and sarcosine oxidase | | | ENSRNOG00000008798 | -0.633 | 0.06718395 |
| Plod1 | procollagen-lysine, 2-oxoglutarate 5-dioxygenase 1 | | | ENSRNOG00000007763 | -0.297 | 0.02647666 |
| Prdm2 | PR/SET domain 2 | | | ENSRNOG00000033522 | -0.232 | 0.16534972 |
| Setd1b | SET domain containing 1B, histone lysine methyltransferase | | | ENSRNOG00000001337 | -0.351 | 0.0825747 |
| Setd2 | SET domain containing 2, histone lysine methyltransferase | | | ENSRNOG00000020915 | -0.176 | 0.16976141 |
| Setdb1 | SET domain bifurcated histone lysine methyltransferase 1 | | | ENSRNOG00000021143 | -0.193 | 0.18525465 |
| Setdb2 | SET domain bifurcated histone lysine methyltransferase 2 | | | ENSRNOG00000021680 | -0.652 | 3.51E-05 |
| **Fatty acid degradation** | | | |  |  |  |
| **Acronym** | **Gene description** | | | **ENSEMBL Reference** | **log2FC** | **Adjusted P** |
| Acaa2 | acetyl-CoA acyltransferase 2 | | | ENSRNOG00000013766 | -0.542 | 0.06876866 |
| Acadl | Acyl-CoA dehydrogenase, long chain | | | ENSRNOG00000012966 | -0.319 | 0.01960642 |
| Acadm | Acyl-CoA dehydrogenase, medium chain | | | ENSRNOG00000009845 | -0.624 | 0.02371996 |
| Acadsb | acyl-CoA dehydrogenase, short/branched chain | | | ENSRNOG00000020624 | -0.416 | 0.08886061 |
| Acadvl | acyl-CoA dehydrogenase, very long chain | | | ENSRNOG00000018114 | -0.284 | 0.00034697 |
| Acat1 | acetyl-CoA acetyltransferase 1 | | | ENSRNOG00000007862 | -0.651 | 0.03091885 |
| Acox1 | Acyl-CoA oxidase 1 | | | ENSRNOG00000008755 | -0.456 | 0.01350453 |
| Acsl1 | Acyl-CoA synthetase long-chain family member 1 | | | ENSRNOG00000010633 | -0.469 | 0.12162677 |
| Acsl4 | Acyl-CoA synthetase long-chain family member 4 | | | ENSRNOG00000019180 | -0.635 | 0.00086761 |
| Aldh2 | aldehyde dehydrogenase 2 family member | | | ENSRNOG00000001344 | -0.297 | 0.0243679 |
| Cpt1a | Carnitine palmitoyltransferase 1A | | | ENSRNOG00000014254 | -0.397 | 0.11355333 |
| Cpt1b | Carnitine palmitoyltransferase 1B | | | ENSRNOG00000010438 | -1.607 | 1.64E-18 |
| Cpt2 | Carnitine palmitoyltransferase 2 | | | ENSRNOG00000012443 | -0.727 | 0.00032609 |
| Cyp4a1 | Cytochrome P450, family 4, subfamily a, polypeptide 1 | | | ENSRNOG00000009597 | -0.812 | 0.00463922 |
| Cyp4a2 | Cytochrome P450, family 4, subfamily a, polypeptide 2 | | | ENSRNOG00000030154 | -0.885 | 0.01482327 |
| Eci1 | enoyl-CoA delta isomerase 1 | | | ENSRNOG00000008843 | -0.873 | 0.00342199 |
| Ehhadh | Enoyl-CoA hydratase and 3-hydroxyacyl CoA dehydrogenase | | | ENSRNOG00000001770 | -1.317 | 4.61E-05 |
| Hadh | hydroxyacyl-CoA dehydrogenase | | | ENSRNOG00000010697 | -0.586 | 0.04364221 |
| Hadha | hydroxyacyl-CoA dehydrogenase trifunctional multienzyme complex subunit α | | | ENSRNOG00000024629 | -0.467 | 0.00285055 |
| Hadhb | hydroxyacyl-CoA dehydrogenase trifunctional multienzyme complex subunit β | | | ENSRNOG00000010800 | -0.659 | 6.28E-05 |
| **Glycerophospholipid metabolism** | | | | |  |  |
| **Acronym** | **Gene description** | | | **ENSEMBL Reference** | **log2FC** | **Adjusted P** |
| Agpat3 | 1-acylglycerol-3-phosphate O-acyltransferase 3 | | | ENSRNOG00000001205 | -0.727 | 8.87E-05 |
| Cds1 | CDP-diacylglycerol synthase 1 | | | ENSRNOG00000002142 | -0.442 | 0.00104103 |
| Cds2 | CDP-diacylglycerol synthase 2 | | | ENSRNOG00000021265 | -0.761 | 5.67E-07 |
| Cept1 | choline/ethanolamine phosphotransferase 1 | | | ENSRNOG00000017723 | -0.512 | 1.91E-06 |
| Chka | choline kinase alpha | | | ENSRNOG00000016791 | -2.512 | 9.80E-70 |
| Chkb | choline kinase beta | | | ENSRNOG00000011404 | -0.951 | 1.84E-17 |
| Dgkd | diacylglycerol kinase, delta | | | ENSRNOG00000023238 | -0.457 | 0.0175824 |
| Gnpat | glyceronephosphate O-acyltransferase | | | ENSRNOG00000019205 | -0.377 | 0.00240699 |
| Gpam | glycerol-3-phosphate acyltransferase, mitochondrial | | | ENSRNOG00000015124 | -2.142 | 2.32E-22 |
| Gpd2 | glycerol-3-phosphate dehydrogenase 2 | | | ENSRNOG00000033824 | -1.699 | 1.44E-25 |
| Lpcat3 | lysophosphatidylcholine acyltransferase 3 | | | ENSRNOG00000012269 | -0.478 | 0.00436203 |
| Lypla2 | lysophospholipase 2 | | | ENSRNOG00000010067 | -0.307 | 0.00802985 |
| Pnpla6 | patatin-like phospholipase domain containing 6 | | | ENSRNOG00000000977 | -0.362 | 0.01746959 |
| Selenoi | selenoprotein I | | | ENSRNOG00000059295 | -1.295 | 1.93E-23 |
| **Fatty acid metabolism** | | |  |  |  |  |
| **Acronym** | **Gene description** | | | **ENSEMBL Reference** | **log2FC** | **Adjusted P** |
| Acaa2 | acetyl-CoA acyltransferase 2 | | | ENSRNOG00000013766 | -0.542 | 0.06876866 |
| Acadl | Acyl-CoA dehydrogenase, long chain | | | ENSRNOG00000012966 | -0.319 | 0.01960642 |
| Acadm | Acyl-CoA dehydrogenase, medium chain | | | ENSRNOG00000009845 | -0.624 | 0.02371996 |
| Acadsb | acyl-CoA dehydrogenase, short/branched chain | | | ENSRNOG00000020624 | -0.416 | 0.08886061 |
| Acadvl | acyl-CoA dehydrogenase, very long chain | | | ENSRNOG00000018114 | -0.284 | 0.00034697 |
| Acat1 | acetyl-CoA acetyltransferase 1 | | | ENSRNOG00000007862 | -0.651 | 0.03091885 |
| Acox1 | acyl-CoA oxidase 1 | | | ENSRNOG00000008755 | -0.456 | 0.01350453 |
| Acsl1 | Acyl-CoA synthetase long-chain family member 1 | | | ENSRNOG00000010633 | -0.469 | 0.12162677 |
| Acsl4 | Acyl-CoA synthetase long-chain family member 4 | | | ENSRNOG00000019180 | -0.635 | 0.00086761 |
| Cpt1a | Carnitine palmitoyltransferase 1A | | | ENSRNOG00000014254 | -0.397 | 0.11355333 |
| Cpt1b | Carnitine palmitoyltransferase 1B | | | ENSRNOG00000010438 | -1.607 | 1.64E-18 |
| Cpt2 | Carnitine palmitoyltransferase 2 | | | ENSRNOG00000012443 | -0.727 | 0.00032609 |
| Ehhadh | Enoyl-CoA hydratase and 3-hydroxyacyl CoA dehydrogenase | | | ENSRNOG00000001770 | -1.317 | 4.61E-05 |
| Elovl2 | ELOVL fatty acid elongase 2 | | | ENSRNOG00000014702 | -0.524 | 0.08893108 |
| Elovl5 | ELOVL fatty acid elongase 5 | | | ENSRNOG00000006331 | -1.332 | 8.20E-20 |
| Hacd2 | 3-hydroxyacyl-CoA dehydratase 2 | | | ENSRNOG00000038761 | -0.281 | 0.07960354 |
| Hadh | hydroxyacyl-CoA dehydrogenase | | | ENSRNOG00000010697 | -0.586 | 0.04364221 |
| Hadha | hydroxyacyl-CoA dehydrogenase trifunctional multienzyme complex subunit α | | | ENSRNOG00000024629 | -0.467 | 0.00285055 |
| Hadhb | hydroxyacyl-CoA dehydrogenase trifunctional multienzyme complex subunit β | | | ENSRNOG00000010800 | -0.659 | 6.28E-05 |
| Hsd17b12 | hydroxysteroid (17-beta) dehydrogenase 12 | | | ENSRNOG00000009630 | -0.377 | 0.01244285 |
| Mcat | malonyl-CoA-acyl carrier protein transacylase | | | ENSRNOG00000010539 | -0.318 | 0.00901845 |
| Mecr | mitochondrial trans-2-enoyl-CoA reductase | | | ENSRNOG00000028047 | -0.638 | 6.28E-06 |
| Oxsm | 3-oxoacyl-ACP synthase, mitochondrial | | | ENSRNOG00000005993 | -0.265 | 0.05729613 |
| Pecr | peroxisomal trans-2-enoyl-CoA reductase | | | ENSRNOG00000055295 | -0.514 | 1.59E-05 |

| **Hepatitis C** |  |  |  |  |
| --- | --- | --- | --- | --- |
| **Acronym** | **Gene description** | **ENSEMBL Reference** | **log2FC** | **Adjusted P** |
| Braf | B-Raf proto-oncogene, serine/threonine kinase | ENSRNOG00000010957 | -0.473 | 0.00169482 |
| Cdkn1a | cyclin-dependent kinase inhibitor 1A | ENSRNOG00000000521 | -2.880 | 4.69E-18 |
| Cldn1 | claudin 1 | ENSRNOG00000001926 | -3.148 | 4.35E-172 |
| Cldn2 | claudin 2 | ENSRNOG00000054495 | -0.588 | 9.57E-06 |
| Cldn14 | claudin 14 | ENSRNOG00000001691 | -0.783 | 0.00039772 |
| Ifit1 | interferon-induced protein with tetratricopeptide repeats 1 | ENSRNOG00000019050 | -1.160 | 0.00028516 |
| Ikbkg | inhibitor of nuclear factor kappa B kinase regulatory subunit gamma | ENSRNOG00000060936 | -0.793 | 5.83E-05 |
| Mapk1 | mitogen activated protein kinase 1 | ENSRNOG00000001849 | -0.350 | 0.00053203 |
| Pik3cb | phosphatidylinositol-4,5-bisphosphate 3-kinase, catalytic subunit beta | ENSRNOG00000016384 | -0.547 | 0.00016849 |
| Ppara | Peroxisome proliferator activated receptor alpha | ENSRNOG00000021463 | -0.874 | 3.58E-05 |
| Psme3 | proteasome activator subunit 3 | ENSRNOG00000051344 | -0.259 | 0.00236065 |
