## Supplementary Table 5 for "Improved Glucose Homeostasis Following Vertical Sleeve Gastrectomy is Associated with Alternate Wiring of the Liver Molecular Clock in a Rat Model of Spontaneously-Occurring Type 2 Diabetes"

**Supplementary Table 5.** Over representation analysis of KEGG biological pathways in response to vertical sleeve gastrectomy in the diabetic Goto-Kakizaki (GK) rat. P-values were calculated following 1000 permutations. Details of rno pathways can be found at [www.genome.jp/kegg](http://www.genome.jp/kegg/). The number of genes considered in the statistical analysis in indicated. FDR, False Discovery Rate.

| **Gene Set** | **Description** | **pValue** | **FDR** | **nb genes** |
| --- | --- | --- | --- | --- |
| rno04064 | NF-kappa B signaling pathway | 4.27 x 10^-11^ | 1.37 x 10^-8^ | 42 |
| rno04670 | Leukocyte transendothelial migration | 2.12 x 10^-8^ | 3.40 x 10^-6^ | 42 |
| rno04380 | Osteoclast differentiation | 3.37 x 10^-8^ | 3.61 x 10^-6^ | 45 |
| rno04062 | Chemokine signaling pathway | 1.39 x 10^-6^ | 1.12 x 10^-4^ | 53 |
| rno05150 | Staphylococcus aureus infection | 3.59 x 10^-6^ | 2.31 x 10^-4^ | 24 |
| rno04650 | Natural killer cell mediated cytotoxicity | 6.98 x 10^-6^ | 3.73 x 10^-4^ | 33 |
| rno04662 | B cell receptor signaling pathway | 3.13 x 10^-5^ | 0.00144 | 25 |
| rno04611 | Platelet activation | 3.66 x 10^-5^ | 0.00147 | 38 |
| rno04710 | Circadian rhythm | 5.21 x 10^-5^ | 0.00182 | 14 |
| rno05202 | Transcriptional misregulation in cancer | 5.66 x 10^-5^ | 0.00182 | 49 |
| rno04659 | Th17 cell differentiation | 6.46 x 10^-5^ | 0.00188 | 33 |
| rno04060 | Cytokine-cytokine receptor interaction | 1.00 x 10^-4^ | 0.00268 | 65 |
| rno04666 | Fc gamma R-mediated phagocytosis | 1.09 x 10^-4^ | 0.00268 | 28 |
| rno04621 | NOD-like receptor signaling pathway | 1.50 x 10^-4^ | 0.00318 | 44 |
| rno01100 | Metabolic pathways | 1.52 x 10^-4^ | 0.00318 | 259 |
| rno03320 | PPAR signaling pathway | 1.64 x 10^-4^ | 0.00318 | 26 |
| rno04625 | C-type lectin receptor signaling pathway | 1.69 x 10^-4^ | 0.00318 | 33 |
| rno04015 | Rap1 signaling pathway | 1.97 x 10^-4^ | 0.00345 | 53 |
| rno04610 | Complement and coagulation cascades | 2.04 x 10^-4^ | 0.00345 | 26 |
| rno04668 | TNF signaling pathway | 2.29 x 10^-4^ | 0.00367 | 32 |
| rno05200 | Pathways in cancer | 2.51 x 10^-4^ | 0.00384 | 112 |
| rno00071 | Fatty acid degradation | 3.95 x 10^-4^ | 0.00537 | 17 |
| rno00620 | Pyruvate metabolism | 4.02 x 10^-4^ | 0.00537 | 15 |
| rno05340 | Primary immunodeficiency | 4.02 x 10^-4^ | 0.00537 | 15 |
| rno05220 | Chronic myeloid leukemia | 5.86 x 10^-4^ | 0.00752 | 24 |
| rno05216 | Thyroid cancer | 7.56 x 10^-4^ | 0.00917 | 14 |
| rno05134 | Legionellosis | 7.72 x 10^-4^ | 0.00917 | 19 |
| rno04810 | Regulation of actin cytoskeleton | 8.35 x 10^-4^ | 0.00925 | 52 |
| rno01212 | Fatty acid metabolism | 8.36 x 10^-4^ | 0.00925 | 18 |
| rno04072 | Phospholipase D signaling pathway | 9.21 x 10^-4^ | 0.00986 | 38 |
| rno05169 | Epstein-Barr virus infection | 9.54 x 10^-4^ | 0.00988 | 55 |
| rno00062 | Fatty acid elongation | 0.00104 | 0.01039 | 13 |
| rno04640 | Hematopoietic cell lineage | 0.00123 | 0.01200 | 27 |
| rno04933 | AGE-RAGE signaling pathway in diabetic complications | 0.00128 | 0.01205 | 28 |
| rno05143 | African trypanosomiasis | 0.00138 | 0.01266 | 14 |
| rno00520 | Amino sugar and nucleotide sugar metabolism | 0.00160 | 0.01391 | 16 |
| rno05140 | Leishmaniasis | 0.00163 | 0.01391 | 22 |
| rno04658 | Th1 and Th2 cell differentiation | 0.00166 | 0.01391 | 26 |
| rno05223 | Non-small cell lung cancer | 0.00169 | 0.01391 | 20 |
| rno05152 | Tuberculosis | 0.00178 | 0.01429 | 44 |
| rno00640 | Propanoate metabolism | 0.00195 | 0.01529 | 12 |
| rno05146 | Amoebiasis | 0.00235 | 0.01781 | 27 |
| rno05142 | Chagas disease (American trypanosomiasis) | 0.00239 | 0.01781 | 28 |
| rno01200 | Carbon metabolism | 0.00277 | 0.02006 | 32 |
| rno04970 | Salivary secretion | 0.00281 | 0.02006 | 22 |
| rno05161 | Hepatitis B | 0.00309 | 0.02155 | 34 |
| rno05144 | Malaria | 0.00319 | 0.02179 | 18 |
| rno05020 | Prion diseases | 0.00352 | 0.02355 | 12 |
| rno05221 | Acute myeloid leukemia | 0.00414 | 0.02713 | 19 |
| rno02010 | ABC transporters | 0.00447 | 0.02859 | 15 |
