## Supplementary Table 6 for "Improved Glucose Homeostasis Following Vertical Sleeve Gastrectomy is Associated with Alternate Wiring of the Liver Molecular Clock in a Rat Model of Spontaneously-Occurring Type 2 Diabetes"

**Supplementary Table 6.** Over representation analysis of Reactome pathways in response to vertical sleeve gastrectomy in the diabetic Goto-Kakizaki (GK) rat. Details of R-RNO pathways can be found at https://reactome.org/. FDR, False Discovery Rate

| **Gene Set** | **Description** | **pValue** | **FDR** | **nb genes** |
| --- | --- | --- | --- | --- |
| R-RNO-168256 | Immune System | 2.44 x 10^-12^ | 3.11 x 10^-9^ | 302 |
| R-RNO-168249 | Innate Immune System | 4.76 x 10^-11^ | 3.04 x 10^-8^ | 187 |
| R-RNO-6798695 | Neutrophil degranulation | 1.02 x 10^-8^ | 4.36 x 10^-6^ | 109 |
| R-RNO-556833 | Metabolism of lipids | 3.02 x 10^-7^ | 9.65 x 10^-5^ | 121 |
| R-RNO-1430728 | Metabolism | 6.68 x 10^-7^ | 1.71 x 10^-4^ | 292 |
| R-RNO-8978868 | Fatty acid metabolism | 1.03 x 10^-6^ | 2.19 x 10^-4^ | 45 |
| R-RNO-449147 | Signaling by Interleukins | 1.30 x 10^-5^ | 0.0021 | 65 |
| R-RNO-193368 | Synthesis of bile acids and bile salts via 7alpha-hydroxycholesterol | 1.33 x 10^-5^ | 0.0021 | 12 |
| R-RNO-1280215 | Cytokine Signaling in Immune system | 2.28 x 10^-5^ | 0.0030 | 81 |
| R-RNO-2162123 | Synthesis of Prostaglandins (PG) and Thromboxanes (TX) | 2.43 x 10^-5^ | 0.0030 | 11 |
| R-RNO-2142753 | Arachidonic acid metabolism | 2.59 x 10^-5^ | 0.0030 | 19 |
| R-RNO-193775 | Synthesis of bile acids and bile salts via 24-hydroxycholesterol | 3.34 x 10^-5^ | 0.0036 | 9 |
| R-RNO-192105 | Synthesis of bile acids and bile salts | 4.17 x 10^-5^ | 0.0041 | 14 |
| R-RNO-194068 | Bile acid and bile salt metabolism | 5.96 x 10^-5^ | 0.0054 | 17 |
| R-RNO-193807 | Synthesis of bile acids and bile salts via 27-hydroxycholesterol | 7.99 x 10^-5^ | 0.0068 | 9 |
| R-RNO-194840 | Rho GTPase cycle | 1.19 x 10^-4^ | 0.0092 | 31 |
| R-RNO-1483257 | Phospholipid metabolism | 1.23 x 10^-4^ | 0.0092 | 44 |
| R-RNO-2454202 | Fc epsilon receptor (FCERI) signaling | 2.38 x 10^-4^ | 0.0158 | 30 |
| R-RNO-983705 | Signaling by the B Cell Receptor (BCR) | 2.44 x 10^-4^ | 0.0158 | 26 |
| R-RNO-9020558 | Interleukin-2 signaling | 2.48 x 10^-4^ | 0.0158 | 7 |
| R-RNO-446652 | Interleukin-1 family signaling | 3.55 x 10^-4^ | 0.0195 | 30 |
| R-RNO-202403 | TCR signaling | 3.59 x 10^-4^ | 0.0195 | 27 |
| R-RNO-9020702 | Interleukin-1 signaling | 3.59 x 10^-4^ | 0.0195 | 27 |
| R-RNO-1280218 | Adaptive Immune System | 3.66 x 10^-4^ | 0.0195 | 121 |
| R-RNO-1483206 | Glycerophospholipid biosynthesis | 3.98 x 10^-4^ | 0.0203 | 31 |
| R-RNO-202424 | Downstream TCR signaling | 4.26 x 10^-4^ | 0.0209 | 24 |
| R-RNO-389948 | PD-1 signaling | 6.04 x 10^-4^ | 0.0285 | 9 |
| R-RNO-5365859 | RA biosynthesis pathway | 7.28 x 10^-4^ | 0.0332 | 11 |
| R-RNO-422475 | Axon guidance | 7.77 x 10^-4^ | 0.0342 | 54 |
| R-RNO-5362517 | Signaling by Retinoic Acid | 8.21 x 10^-4^ | 0.0350 | 15 |
| R-RNO-202433 | Generation of second messenger molecules | 8.79 x 10^-4^ | 0.0362 | 10 |
| R-RNO-187577 | SCF(Skp2)-mediated degradation of p27/p21 | 0.0012 | 0.0450 | 17 |
| R-RNO-109582 | Hemostasis | 0.0012 | 0.0451 | 93 |
| R-RNO-202430 | Translocation of ZAP-70 to Immunological synapse | 0.0012 | 0.0451 | 7 |
| R-RNO-512988 | Interleukin-3, Interleukin-5 and GM-CSF signaling | 0.0013 | 0.0460 | 12 |
| R-RNO-2871837 | FCERI mediated NF-kB activation | 0.0013 | 0.0460 | 20 |
| R-RNO-5607764 | CLEC7A (Dectin-1) signaling | 0.0013 | 0.0460 | 23 |
| R-RNO-69202 | Cyclin E associated events during G1/S transition | 0.0014 | 0.0477 | 18 |
| R-RNO-8878166 | Transcriptional regulation by RUNX2 | 0.0015 | 0.0477 | 17 |
| R-RNO-8939902 | Regulation of RUNX2 expression and activity | 0.0015 | 0.0477 | 15 |
| R-RNO-202733 | Cell surface interactions at the vascular wall | 0.0015 | 0.0477 | 26 |
| R-RNO-2871809 | FCERI mediated Ca+2 mobilization | 0.0017 | 0.0509 | 9 |
| R-RNO-69656 | Cyclin A:Cdk2-associated events at S phase entry | 0.0018 | 0.0538 | 18 |
| R-RNO-2142691 | Synthesis of Leukotrienes (LT) and Eoxins (EX) | 0.0022 | 0.0634 | 7 |
| R-RNO-202427 | Phosphorylation of CD3 and TCR zeta chains | 0.0022 | 0.0634 | 7 |
| R-RNO-76002 | Platelet activation, signaling and aggregation | 0.0024 | 0.0636 | 50 |
| R-RNO-114604 | GPVI-mediated activation cascade | 0.0024 | 0.0636 | 11 |
| R-RNO-166016 | Toll Like Receptor 4 (TLR4) Cascade | 0.0024 | 0.0636 | 23 |
| R-RNO-5621481 | C-type lectin receptors (CLRs) | 0.0031 | 0.0787 | 26 |
| R-RNO-168898 | Toll-like Receptor Cascades | 0.0031 | 0.0787 | 29 |
