## Supplementary Table 3 for "Improved Glucose Homeostasis Following Vertical Sleeve Gastrectomy is Associated with Alternate Wiring of the Liver Molecular Clock in a Rat Model of Spontaneously-Occurring Type 2 Diabetes"

**Supplementary Table 3.** Biological pathways significantly affected by vertical sleeve gastrectomy in the diabetic Goto-Kakizaki (GK) rat. Gene set enrichment analysis (GSEA) was used to identify significantly enriched or downregulated KEGG pathways in the liver transcriptome of gastrectomised GK rats. P-values were calculated following 1000 permutations. Details of pathways in Rattus Novergicus (rno) can be found at [www.genome.jp/kegg](http://www.genome.jp/kegg/). Size is the total number of genes in the pathway and count is the number of sequenced genes considered in the statistical analysis. NES, normalised enrichment score. FDR, False Discovery Rate.

| **rno** | **Pathway name** | **NES** | **P value** | **FDR** | **Size** | **Count** |
| --- | --- | --- | --- | --- | --- | --- |
| 04940 | Type I diabetes mellitus | 1.851 | 0.004 | 0.222 | 40 | 17 |
| 04610 | Complement and coagulation cascades | 1.836 | 0.017 | 0.208 | 72 | 35 |
| 04964 | Proximal tubule bicarbonate reclamation | 1.809 | 0.006 | 0.205 | 18 | 6 |
| 05320 | Autoimmune thyroid disease | 1.806 | 0.005 | 0.189 | 34 | 16 |
| 04713 | Circadian entrainment | 1.805 | 0.021 | 0.175 | 66 | 17 |
| 04380 | Osteoclast differentiation | 1.770 | 0.018 | 0.181 | 108 | 45 |
| 05323 | Rheumatoid arthritis | 1.770 | 0.026 | 0.169 | 57 | 23 |
| 05322 | Systemic lupus erythematosus | 1.759 | 0.007 | 0.164 | 51 | 26 |
| 05310 | Asthma | 1.733 | 0.006 | 0.169 | 12 | 9 |
| 05340 | Primary immunodeficiency | 1.711 | 0.019 | 0.171 | 26 | 21 |
| 04650 | Natural killer cell mediated cytotoxicity | 1.708 | 0.019 | 0.164 | 77 | 33 |
| 05143 | African trypanosomiasis | 1.688 | 0.011 | 0.166 | 25 | 13 |
| 04062 | Chemokine signaling | 1.681 | 0.025 | 0.162 | 146 | 63 |
| 04623 | Cytosolic DNA-sensing | 1.670 | 0.009 | 0.161 | 42 | 19 |
| 04672 | Intestinal immune network for IgA production | 1.651 | 0.025 | 0.165 | 28 | 18 |
| 05152 | Tuberculosis | 1.614 | 0.038 | 0.181 | 130 | 43 |
| 05133 | Pertussis | 1.558 | 0.026 | 0.222 | 59 | 36 |
| 05142 | Chagas disease (American trypanosomiasis) | 1.554 | 0.027 | 0.217 | 88 | 32 |
| 04625 | C-type lectin receptor signaling | 1.546 | 0.033 | 0.218 | 91 | 31 |
| 00440 | Phosphonate and phosphinate metabolism | -1.548 | 0.019 | 0.443 | 5 | 2 |
| 00604 | Glycosphingolipid biosynthesis | -1.584 | 0.033 | 0.432 | 13 | 2 |
| 04950 | Maturity onset diabetes of the young | -1.601 | 0.042 | 0.433 | 15 | 5 |
| 04670 | Leukocyte transendothelial migration | -1.610 | 0.027 | 0.447 | 92 | 8 |
| 04146 | Peroxisome | -1.616 | 0.044 | 0.475 | 76 | 28 |
| 00970 | Aminoacyl-tRNA biosynthesis | -1.649 | 0.032 | 0.453 | 43 | 25 |
| 00310 | Lysine degradation | -1.671 | 0.033 | 0.454 | 54 | 23 |
| 04728 | Dopaminergic synapse | -1.673 | 0.046 | 0.502 | 94 | 2 |
| 00071 | Fatty acid degradation | -1.763 | 0.030 | 0.385 | 39 | 20 |
| 01040 | Biosynthesis of unsaturated fatty acids | -1.799 | 0.013 | 0.370 | 24 | 9 |
| 00564 | Glycerophospholipid metabolism | -1.807 | 0.029 | 0.416 | 78 | 14 |
| 01212 | Fatty acid metabolism | -1.818 | 0.016 | 0.473 | 45 | 24 |
| 05160 | Hepatitis C | -2.031 | <10^-4^ | 0.259 | 101 | 11 |
| 03320 | PPAR signaling | -2.074 | 0.004 | 0.298 | 62 | 21 |
