## Supplementary figures and images for "Improved Glucose Homeostasis Following Vertical Sleeve Gastrectomy is Associated with Alternate Wiring of the Liver Molecular Clock in a Rat Model of Spontaneously-Occurring Type 2 Diabetes"

### Supplementary Figure 1

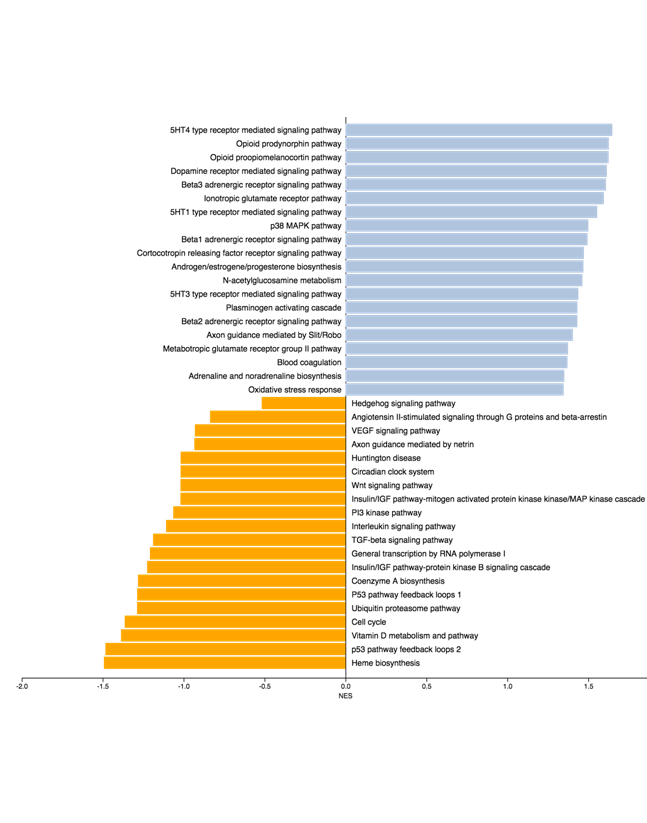

### Supplementary Figure 2

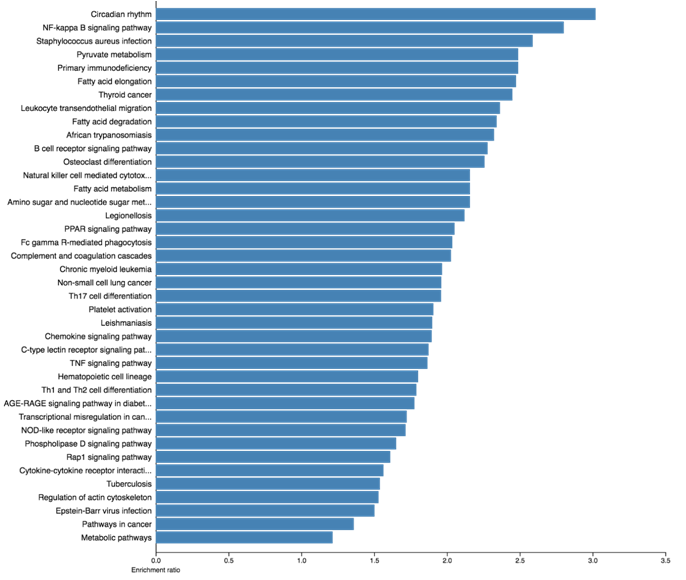

### Supplementary Figure 3

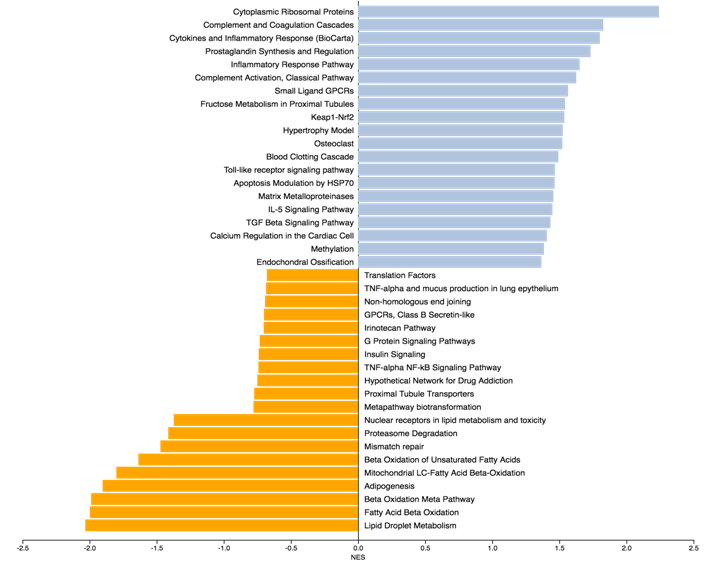

### Supplementary Figure 4

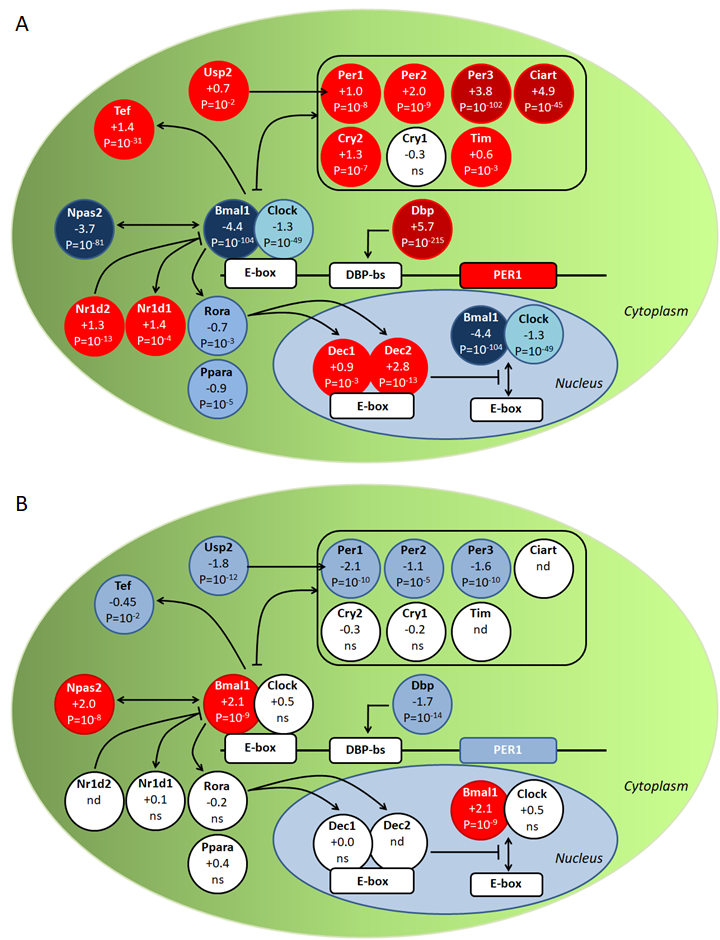
