## Supplementary Table 1 for "Improved Glucose Homeostasis Following Vertical Sleeve Gastrectomy is Associated with Alternate Wiring of the Liver Molecular Clock in a Rat Model of Spontaneously-Occurring Type 2 Diabetes"

Supplementary Table 1. Oligonucleotides used in qPCR experiments in GK rats.

| **Gene** | **Forward (5’ – 3’)** | **Reverse (5’ – 3’)** |
| --- | --- | --- |
| ***Arntl*** | TCAGGCCTACACAGCTACAC | AGTTCTCAAGCTTCGGTCCA |
| ***Ciart*** | GACGGTGAAGGAGCTGGATA | GGGAGAGAGAGGAGGAGGAA |
| ***Clock*** | CCATGGTCAAGGGCTACAGA | CACCCCACTCTGAATCTGGT |
| ***Cry2*** | GTCCTTCCACCTCCTTGACA | TGCAACCATCTCTTCCCCAT |
| ***F4/80/Adgre1*** | GCCCTTCCAACTCATCATGT | AGGGAATCCTTTTGCATGTG |
| ***Gata3*** | GAGAGCAGGGACTTCCTGTG | CATCATGCACCTTTTTGCAC |
| ***Il1b*** | CTGGTACATCAGCACCTCTCAA | GAGACTGCCCATTCTCGACAA |
| ***Il10*** | CTGCAGGACTTTAAGGGTTACTTG | TTCTCACAGGGGAGAAATCG |
| ***Mcp1/Ccl2*** | CTGGACCAGAACCAAGTGAGATCA | GTGCTTGAGGTGGTTGTGGAAA |
| ***MhcII/Rt1bb*** | CCCTCCAGCGGTCAATGT | TGACACGCCTTTGGTGACA |
| ***Nfκb/Rela*** | CTGGCCATGGACGATCTGTTT | CCCTCGCACTTGTAACGGAAA |
| ***Npas2*** | GCAAAGGGAAGTCGTGTTGT | GCTCCTGTCTCCTTTCCACT |
| ***Nr1h4 (Fxr)*** | TACCATTACAACGCGCTCAC | GCCCCCGTTCTTACACTTG |
| ***Per1*** | CCTGTTTTGTCCTCCACTGC | GCATCAGTGTCATCAGCCAG |
| ***Per2*** | ATCTCCAGGCGGTCTTGAAA | ACACACCCTGTTACGTCGAT |
| ***Per3*** | CGCAGAAGGAAGAGCAGAAC | TGTGCTTCCGTCTCTCACTT |
| ***Ppara*** | CAATGGCTTCATCACCCGAG | ATCCCCTCCTGCAACTTCTC |
| ***Rora*** | CACACACCCAACATGAAGCA | TCTCCCTCCCTCTTCATCCA |
| ***Srebf1*** | CCGTTTCTTCGTGGATGG | CACAGAATAGTCGGGTCACCT |
| ***Tgfb1*** | GAGCCCGAGGCGGACTACTA | CCCGAATGTCTGACGTATTGAAGA |
| ***Tnfa*** | TGAACTTCGGGGTGATCG | GGGCTTGTCACTCGAGTTTT |
